# Liquid-liquid phase separation of tau driven by hydrophobic interaction facilitates fibrillization of tau

**DOI:** 10.1101/2020.08.05.237966

**Authors:** Yanxian Lin, Yann Fichou, Andrew P. Longhini, Luana C. Llanes, Yinson Yin, Guillermo C. Bazan, Kenneth S. Kosik, Songi Han

## Abstract

Amyloid aggregation of tau protein is implicated in neurodegenerative diseases, yet its facilitating factors are poorly understood. Recently, tau has been shown to undergo liquid liquid phase separation (LLPS) both *in vivo* and *in vitro*. LLPS was shown to facilitate tau amyloid aggregation in certain cases, while independent of aggregation in other cases. It is therefore important to understand the differentiating properties that resolve this apparent conflict. We report on a model system of hydrophobically driven LLPS induced by high salt concentration (LLPS-HS), and compare it to electrostatically driven LLPS represented by tau-RNA/heparin complex coacervation (LLPS-ED). We show that LLPS-HS promotes tau protein dehydration, undergoes maturation and directly leads to canonical tau fibrils, while LLPS-ED is reversible, remains hydrated and does not promote amyloid aggregation. We show that the nature of the interaction driving tau condensation is the differentiating factor between aggregation-prone and aggregation-independent LLPS.

## Introduction

Tau is a protein mainly present in the central nervous system that binds and stabilizes microtubules in neurons [1]–[3]. Under healthy conditions, tau is soluble across a wide range of temperatures, protein concentrations, pH and ionic strengths of the buffer, while under pathologic conditions, tau undergoes amyloid aggregation that yields insoluble, irreversible and cross-β rich fibrils. While the functions of tau are not fully understood, in the healthy state tau is tightly bound to microtubules [4] and the concentration of the unbound tau protein is exceptionally low—on the order of 1∼10 nM ^1^. Under pathological conditions, intracellular tau transforms from its microtubule-bound or soluble state to a fibrillar state. In fact, intracellular deposits of aggregated tau are a diagnostic hallmark of a wide range of neurodegenerative diseases known as tauopathies [8]. The mechanism by which low concentrations of tau is recruited to assemble into tangles remains undetermined. Hence, factors that drive assembly and condensation, or generally mechanisms that increase the intracellular local concentration of tau are of great interest to understand this important pathological process.

Liquid-liquid phase separation (LLPS) is a process that is readily driven by multivalent weak interactions between flexible polymers, yielding a polymer-rich phase separated from a polymer-depleted phase [9], [10], the former of which is usually observed as micrometer size fluidic droplets. Here, we use the widest definition of polymers that hence includes intrinsically disordered proteins (IDPs) and ribonucleic acid (RNA). LLPS has been used to describe the formation of membraneless organelles *in vivo* that are proposed to play important roles in mediating cellular functions [11], [12]. The interest in biomolecular condensates in the cellular context has exploded in recent years, fueled by intriguing observations of LLPS of IDPs associated with neurodegenerative diseases, including FUS, TDP-43, hnRNPA1, synuclein [13]–[18] as well as more recently tau [19]–[25]. Despite the ubiquitous occurrence of LLPS with IDPs, the physical principles that govern their formation, phase diagrams or properties are not universal. LLPS of IDPs can be driven by a variety of interaction types and with a wide range of interaction strengths. The driving interaction for association that forms the basis for LLPS typically includes electrostatic charge-charge and dipole-dipole interactions between the backbone, charged/polar side chains and polar solvent molecules, and hydrophobic interactions. The latter involve water-repelling protein or polymer side chains that can be aromatic [26], non-polar [9], [27] or even charged residues [28], [29] that, in the context of the protein or polymer surface, may display less favorable protein-water interactions compared to protein-protein and water-water interactions. Molecular crowding, often modeled by adding polyethylene glycol (PEG), can therefore enhance hydrophobic interactions by removing hydration water from the dense LLPS phase [30]. However, the potentially differential role of electrostatic and hydrophobic interactions for the LLPS of tau have neither been investigated nor dissected to date. Recent work [9], [26], [31]–[35] has highlighted certain interactions as particularly important in driving protein LLPS, in particular π-π and π-cation interaction between charged and aromatic residues [36], [37]. Tau has few aromatic amino acids (∼2% for full length tau 2N4R) or arginine residues (∼3% for 2N4R), consequently little possibility to engage in π-π or π-cation interactions. Nevertheless, tau’s sequence exhibits a rich landscape of chemical properties with polar, charged and hydrophobic residues present throughout the protein. There is a need to study the driving forces of tau LLPS and uncover what role they play in pathology.

LLPS has been shown to promote amyloid aggregation of FUS and hnRNPA1 [13]–[16]. The verdict is not so clear for LLPS of tau. Ambadipudi *et al*. in 2017 [19] and Wegmann *et al*. in 2018 [20] proposed that tau in a LLPS state actively promoted amyloid aggregation, while a recent study from our group [38] showed that tau in LLPS formed by complex coacervation with RNA does not induce or protect from aggregation. Herein, we extensively explore the hypothesis that the nature of the interactions driving tau LLPS is the main factor that influence how LLPS impact amyloid aggregation.

We and others have previously shown that tau undergoes electrostatically driven LLPS (LLPS-ED), either with itself—in a process known as simple coacervation (SC)—or with polyanions—in a process known as complex coacervation (CC) [21], [22], [33], [38]. Meanwhile, tau can undergo LLPS upon addition of high concentration of salt (LLPS-HS)[38]. Concentrated salt (e.g. of NH_4_^+^, K^+^ or Na^+^ along the Hofmeister series) has been shown to exert a “salting-out” effect [39], [40], presumably by dehydrating the protein and promoting entropy-driven hydrophobic interactions among proteins [41], [42]. High concentration of salt including NaCl has been found to induce LLPS of several proteins, including BSA [43], lysozyme [44] and most recently FUS, TDP-43, and Annexin A11 [45]. In this article, we use the condensation of tau induced by adding high concentration of salt (LLPS-HS) as a model for tau LLPS driven by hydrophobic interactions. Using EPR spectroscopy, fluorescence spectroscopy and biochemical assays, we compared the properties of tau in LLPS-HS with that of LLPS-ED formed by tau-tau or tau-RNA to uncover whether there is a cause-effect relationship between the LLPS and aggregation of tau.

## RESULTS

### Tau undergoes LLPS at high salt concentration

The longest human tau isoform (2N4R) is a charged and hydrophilic IDP near physiological conditions, displaying both high solubility and stability. The charge and Hopp-Woods hydrophobicity plots (both are moving-averaged over 25 consecutive residues) show that 2N4R consists of a hydrophilic and negatively charged N-terminal half and a weakly hydrophobic and overall positively charged C-terminal half (**Figure 1**A). The four repeat domains (R1–R4) are responsible for binding microtubules and constitute a large part of the core of amyloid fibrils found *in vivo* and *in vitro* [46]. The goal of our study is to investigate tau LLPS based on hydrophobic interactions separately from those based on electrostatic interactions. To maximize the hydrophobicity while keeping the repeat domains, we studied the LLPS of N-terminal-truncated 2N4R between residues 255-441, referred to as tau187 (**Figure 1**A). To mute potential effects of covalent disulfide bonding, we introduced double site mutations to the two native cysteines, C291S/C322S. Unless stated explicitly, we refer to tau187C291SC322S as tau187, and to 2N4RC291SC322S as 2N4R throughout this work.

**Figure 1.**
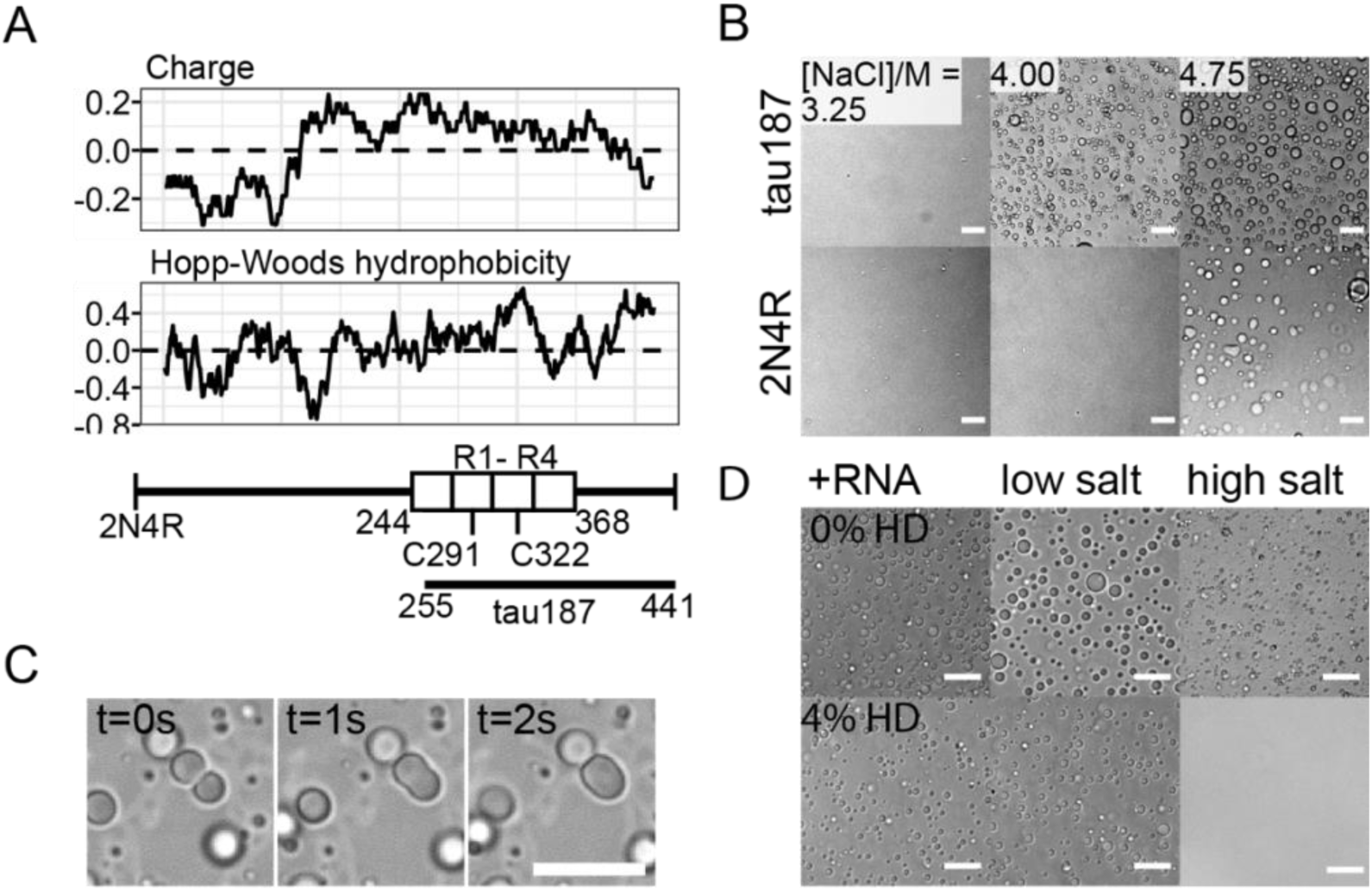
Tau undergoes LLPS at high salt concentration. **A**. Charge and Hopp-Woods hydrophobicity plot of 2N4R. Data points show the average values of consecutive 25 amino acids. Reference values of Glycine are shown as dashed lines. tau187 fragment (residues 255-441), four repeat domains (R1–R4, 244-368) and two native cysteines (C291, C322) are shown. **B**. Microscope images of tau187 and 2N4R at varying [NaCl]. 44 µM tau187 and 20 µM 2N4R were chosen to have same mass concentration. Scale bar length is 25 µm. **C**. Droplet merging of tau LLPS-HS. 450 µM tau187 with 2.2 M NaCl was used. Scale bar length is 50 µm. **D**. Effects of 4 wt% 1,6-hexanediol on different types of tau LLPS. Tau-RNA LLPS (+RNA) was prepared with 20 µM 2N4R and 40 µg/mL polyU RNA; Tau low salt simple coacervation (low salt) was prepared with 20 µM 2N4R at 5 mM NaCl; Tau high salt simple coacervation (high salt) was prepared with 100 µM tau187 with 4.75 M NaCl. Scale bar length is 25 µm.

To eliminate electrostatic interactions, we mixed tau with excess amount of NaCl. Both tau187 and 2N4R samples became turbid and formed abundant droplets when [NaCl] was adjusted to above 4 M and 4.75 M, respectively (**Figure 1**B). After formation, droplets merged within seconds, confirming their fluidity (**Figure 1**C). We refer to this process of high salt concentration-induced LLPS as LLPS-HS, and the resulting droplets as high salt droplets in this article. We furthermore confirmed that both tau187C291S and tau187C291SP301L form LLPS-HS (Figure 1-figure supplement 1), showing that inter-tau disulfide bonding and aggregation-prone mutations do not prevent LLPS-HS.

We have previously shown that tau187 undergoes LLPS-ED, and established a phase diagram using a combination of experiments, theory underlying coacervation and field theoretic simulations (FTS) of LLPS [22]. The LLPS-ED phase (formed either by SC or CC) typically dissolves at a total salt concentration above [NaCl] ∼ 100-200 mM [22], [33], while the LLPS-HS obviously only forms above a much higher salt concentration (3-4 M; **Figure 1**B). The question we ask is whether the properties of the protein tau and the LLPS phase are different between the two forms of LLPS. We start by testing the sensitivity of the dense liquid phase to 1,6-hexanediol (1,6-HD), an amphiphilic small molecule that is known to disrupt weak hydrophobic interactions, [47]–[49] which has been widely used to verify the formation and dissolution of various forms of LLPS, including that of FUS [50] and of tau [20], [51]. We prepared tau LLPS-ED in two ways: by mixing tau187 with RNA by the CC mechanism, or by lowering the ionic strength of the solution containing 2N4R (< 10 mM, referred to as low-salt droplets) that induces SC (**Figure 1**D). When adding 4% 1,6-HD, neither form of tau LLPS-ED, tau-RNA droplets formed by CC or tau low-salt droplets formed by SC, dissolved. In contrast, tau LLPS-HS dissolved completely upon addition of 4% 1,6-HD (**Figure 1**D).

The distinct effect of 1,6-HD demonstrates that the tau molecules are held together in the condensed phase of LLPS by different types of interactions in LLPS-ED versus in LLPS-HS. The observation that the stability of LLPS-ED, whether formed of tau187-RNA CC or of 2N4R SC, is insensitive to 1,6-HD confirmed that the interactions holding tau together in the condensed phase is not weakened by an amphiphilic molecule; this behavior is expected for purely electrostatic associations. Considering that the Debye length at 4 M NaCl is ∼0.1 nm, which is much smaller than the diameter of a water molecule [52], [53], electrostatic interactions between tau molecules are muted in LLPS-HS. Besides electrostatic screening, salts made of Na^+^ and Cl^-^ at these concentrations also exert dehydration induced salting out of proteins following the Hofmeister series. Together with the observation that 1,6-HD dissolves high salt droplets, these results demonstrate that LLPS-HS are predominantly hydrophobically driven. Hydrophobic interactions arise from weaker protein-water compared to water-water and protein-protein interactions, resulting in (i) water to be readily expelled from the protein surface to rather hydrogen bond with other solvent molecules, i.e. other water molecules in the bulk, and (ii) protein molecules to interact via van der Waals interactions. Amphiphilic molecules such as 1,6-HD can restore interactions between protein and water and/or disrupt interaction between associated proteins, disrupting the hydrophobic driving forces for LLPS-HS. These results verify that LLPS-HS is a good model system to study the consequences of hydrophobically driven LLPS on the irreversible process of aggregation.

### LLPS-HS undergoes irreversible maturation

Many examples of protein LLPS have been shown to undergo irreversible maturation [16], [54]. To study the reversibility property of LLPS-HS, we recorded turbidity of high salt droplet samples upon repeated heating-cooling cycles. At 3.5 M NaCl, turbidity at T < 35 °C remains near zero and significantly increases to 0.4 at T > 40 °C, indicating a lower critical saturation temperature behavior (LCST) where LLPS is thermodynamically more favorable at higher temperature (**Figure 2-figure supplement 1**A). When cooled down to low temperature, the original transparent sample remained slightly turbid (**Figure 2-figure supplement 1**A), a second heating-cooling cycle showed further increase of remaining turbidity. In contrast, tau-RNA LLPS-ED formed by CC showed no remaining turbidity after consecutive heating-cooling cycles (**Figure 2-figure supplement 1**B). The same results were found with 2N4R (**Figure 2-figure supplement 1**C). The hysteresis in the build-up of turbidity with repeated heating-cooling cycle implies that LLPS-HS leads to partially irreversible associations between tau, while LLPS-ED formation and melting is reversible.

**Figure 2.**
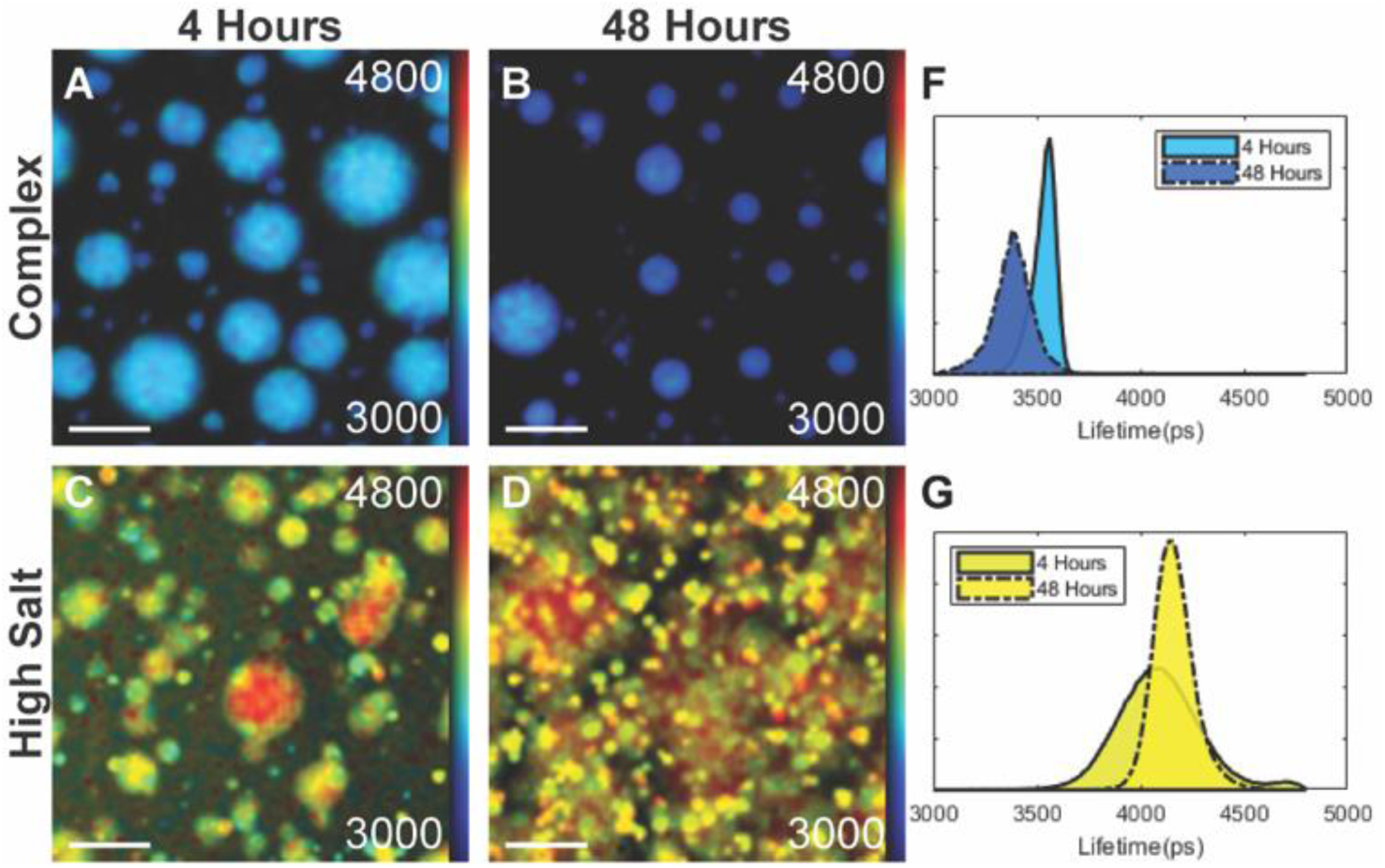
FLIM fits of tau-RNA LLPS-CC (complex), and tau LLPS-high salt (high salt). Fluorescence Lifetime Imaging Measurements were taken at 4 and 48 hours at the same resolution, scale bar = 25 μm. **A-D**, Fluorescent microscope images of tau-RNA droplets and tau high salt droplets before (4 hour) and after incubation (48 hour). Individual pixels were fitted with a 2-component exponential decay using the FLIMfit software tool developed at Imperial College London (Warren SC et al, PLOS ONE 2013), and the data were visualized using the higher lifetime component. The colors in the fluorescent images and corresponding histograms represent a pseudo color heat map that ranges from 3.0 to 4.8 ns, with blue representing low lifetimes and red representing high lifetimes. **F-G**. Histograms of fitted results in A-D. The histograms are normalized so that area under the curve is unity. The color under each histogram corresponds to the pseudo color of the heat map at the histograms max value. Solid line = 4 hours, Dashed line = 48 hours.

To further probe the micro-environment inside droplets, we labeled tau with an in-house synthesized fluorescence molecular rotor derived from a boron-dipyrromethene (BODIPY) core [55] and performed Fluorescence Lifetime Image Measurements (FLIM) over the course of 48 hours (See Materials and Methods for synthesis and microscopy details). Molecular rotors are a class of dyes whose fluorescent lifetimes are sensitive to the local microviscosity of their environment, with a low lifetime indicating a fluid, water-like environment and a high lifetime representing a viscous environment [56]–[58]. Further, within a limited regime, the relationship between the fluorescent lifetime and the microviscosity follows the Förster-Hoffman equation (see calibration between fluorescent lifetime and viscosity in **Figure 2-figure supplement 2**). Unlike viscosity that describes the bulk property of a liquid, microviscosity describes the friction that, in this case, is experienced by a single biomolecule due to its local environment. The molecular rotor was conjugated to 2N4R, as described in Materials and Methods. FLIM images were collected on a confocal microscope and a pixelwise fit of the images binned into histograms. Shortly after triggering LLPS, 2N4R LLPS-HS droplets showed a Gaussian distribution of lifetimes centered around 4.08±0.32 ns (916 cP). In comparison, 2N4R LLPS-ED droplets formed by CC between 2N4R and RNA (**Figure 2**E, F) showed a lifetime centered around 3.55±0.07 ns (195 cP). In other words, 2N4R in LLPS-HS experiences a higher microviscosity than in LLPS-ED. After a 48 hour incubation period, the average fluorescence lifetime of 2N4R in LLPS-HS samples increased from 4.08±0.26 to 4.15±0.13 ns (**Figure 2**F), while that of 2N4R in LLPS-ED decreased from 3.55±0.07 ns to 3.39±0.12 ns (**Figure 2**E, F). Concurrently, amorphous, non-spherical condensates formed within LLPS-HS, while the overall droplet shape of LLPS-ED (of 2N4R-RNA CC) remains spherical (**Figure 2**B, D). The emergence of non-spherical structures within the droplets and the increase in microviscosity sensed by the tau protein indicate that LLPS-HS undergo maturation of the condensate. LLPS-ED on the other hand retains liquid properties that appears to slightly less viscous over time.

### LLPS-HS triggers canonical tau amyloid aggregation

To test the hypothesis that maturation of droplets is associated with amyloid formation in the sample, we incubated high salt droplets with a trace amount of Thioflavin T (ThT). While high salt droplets emerge spontaneously and fully upon preparing samples under droplet-forming conditions, ThT fluorescence starts from baseline and steadily increases over ∼24 hour duration (**Figure 3**A). The increase of ThT fluorescence reflects an increase in cross-β sheet content developing over hours. Subsequent imaging of tau samples—in LLPS-HS conditions after overnight incubation— showed fibril-like species by TEM that are visually similar to tau-heparin fibrils (**Figure 3**B). This supported the hypothesis that LLPS-HS is followed by the amyloid aggregation of tau.

**Figure 3.**
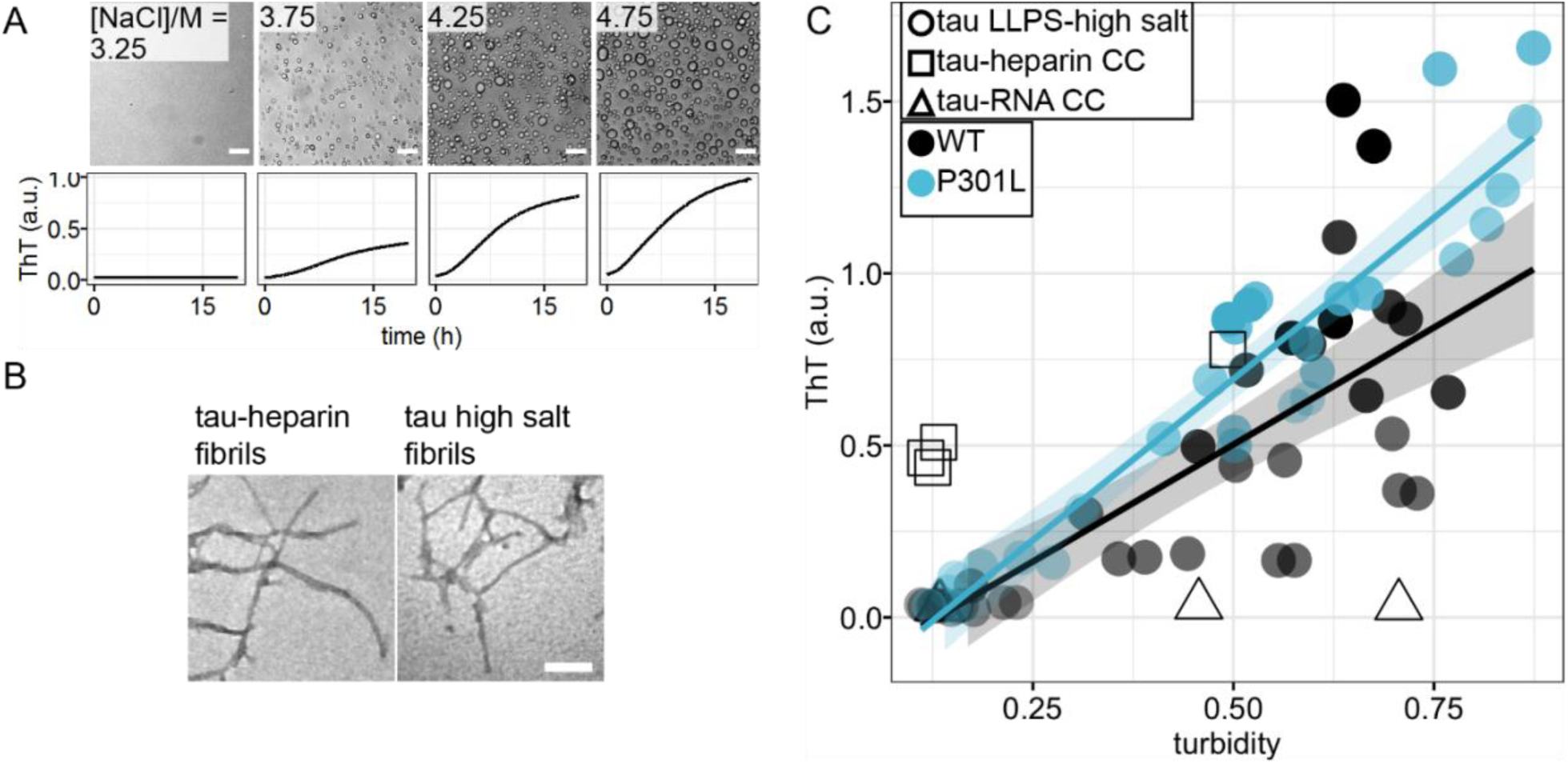
Correlation of LLPS-high salt and amyloid aggregation. **A**. Representative microscope images at t = 0 and ThT fluorescence at room temperature overnight of tau LLPS-high salt at various [NaCl]. 44 µM tau187 was used. **B**. Representative TEM image of tau LLPS-high salt sample after overnight incubation, compared with tau-heparin fibrils. **C**. Correlation of initial turbidity and final ThT fluorescence of tau LLPS-high salt vs LLPS-CC. Solids line and shadow show linear regression and its standard error, respectively. 40 *μM* ± 10% of tau187 was used. Samples come from 5 different batches of proteins.

To show the connection between LLPS-HS and amyloid aggregation, we incubated tau at equal concentration, while varying [NaCl] on a multi-well plate. We recorded bright field microscope images before incubation showing the amount of high salt droplets, after which we monitored ThT fluorescence over time (**Figure 3**A). The abundance of droplets visually observed via bright field microscope imaging immediately after sample preparation (at *t* = 0) strongly correlates with the ThT fluorescence intensity after incubation (*t* = 24 hours). The same results were found with 2N4R (**Figure 3-figure supplement 2**). The initial turbidity (at 500 nm) that quantifies the volume fraction of high salt droplets at *t* = 0 (Figure SI 10 in [38]) was plotted against final ThT fluorescence readings at *t* = 24 hours, representing the relative amount of amyloid aggregates formed (**Figure 3**C). We found a proportional correlation (linear R_2_ of 0.43) between the initial turbidity and the final ThT fluorescence readings (**Figure 3**C). This result implies that LLPS-HS causally triggers amyloid formation. In contrast, electrostatically driven LLPS neither promotes RNA-assisted tau aggregation, nor prevents heparin-induced tau aggregation as shown in **Figure 3**C and **Figure 3-figure supplement 1**, and as previously reported [38]. These results show that the strong correlation between LLPS and amyloid aggregation is unique to hydrophobically-driven LLPS, as represented by LLPS-HS, while LLPS-ED and aggregation are independent processes.

We further studied the effects of the frontotemporal dementia-related mutation P301L of tau on the correlation between LLPS-HS and aggregation. Our data showed that tau187P301L shows an even steeper dependence of the final ThT fluorescence intensity to the initial droplet quantity at *t* = 0 (**Figure 3**C), implying that for a similar initial volume of droplets, tau187P301L generates higher quantities of cross β-sheet structures, resulting in higher ThT fluorescence intensity. Meanwhile, an analysis of the aggregation half time showed that in all examined conditions P301L has significantly shortened the half time (**Figure 6-figure supplement 2**). These results showed that P301L promotes tau aggregation under LLPS-HS conditions, consistent with its broadly known aggregation-promoting properties under various *in vitro* and *in vivo* conditions. Similar results were observed with 2N4R (**Figure 6-figure supplement 1**).

**Figure 4.**
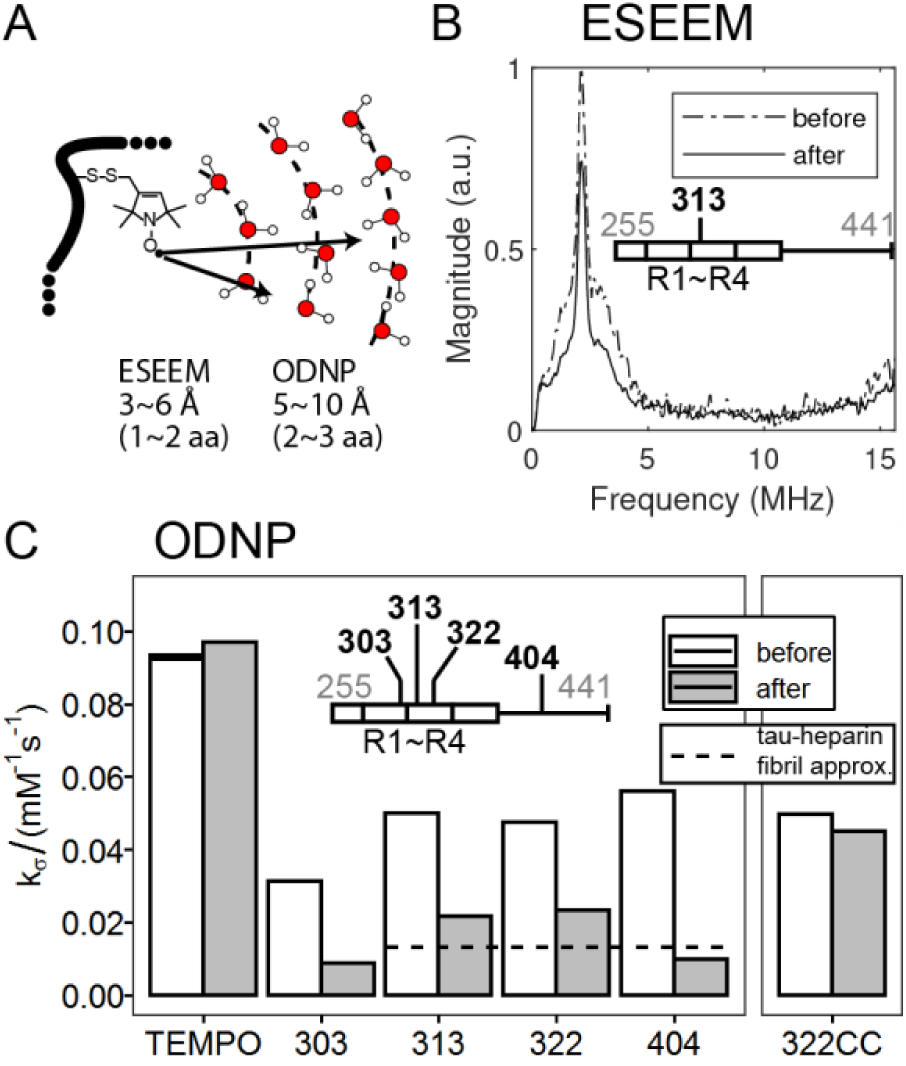
LLPS high salt undergoes dehydration. **A**. Schematic diagram showing sensitive hydration shell in Electron Spin Echo Envelope Modulation (ESEEM) and Overhauser Nuclear Dynamic Polarization (ODNP). **B**. Representative 3 pulse-ESEEM of tau187 at site 313 at solution (before) and upon addition of 3.75 M NaCl (after). 22.5% Ficoll was used as glassing reagent. **C**. Representative ODNP cross-relaxivity parameter *k*_*σ*_ of tau187 at various sites at solution (before) and upon addition of 3.75 M NaCl (after). 250 µM tau was used. For comparison, *k*_*σ*_ of tau187 at site 322 upon tau-RNA complex coacervation was shown (322CC). 80% reduction of *k*_*σ*_ has been reported at site 313, 322 and 404 for tau-heparin fibrils (Pavlova, et al 2016), and is shown as dashed line.

**Figure 5.**
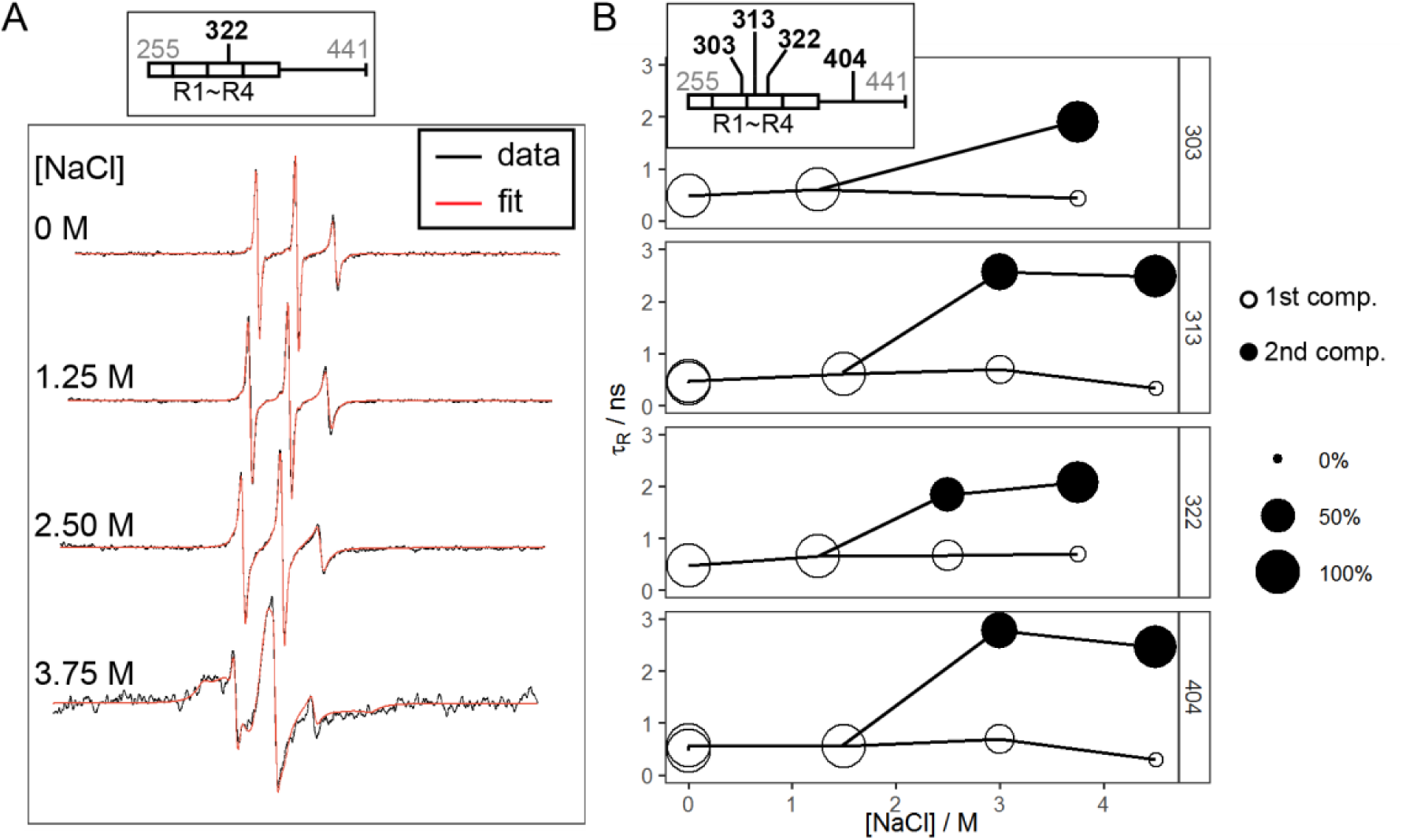
Site specific dynamics of tau187 upon LLPS-high salt. **A**. Representative X-band cwEPR lineshape of tau187 at site 322 at varying [NaCl]. Black lines show the experiment data and red lines show the fit. Spectra are shifted to avoid overlap. **B**. Rotational correlation time, τ_*R*_, of tau187 at site 303, 313, 322 and 404 at varying [NaCl]. Data at low [NaCl] were fit with 1 component with the y-axis showing the τ_*R*_. Data at high [NaCl] were fitted with 2 components with the area of the disk proportional to the percentage of each component.

**Figure 6.**
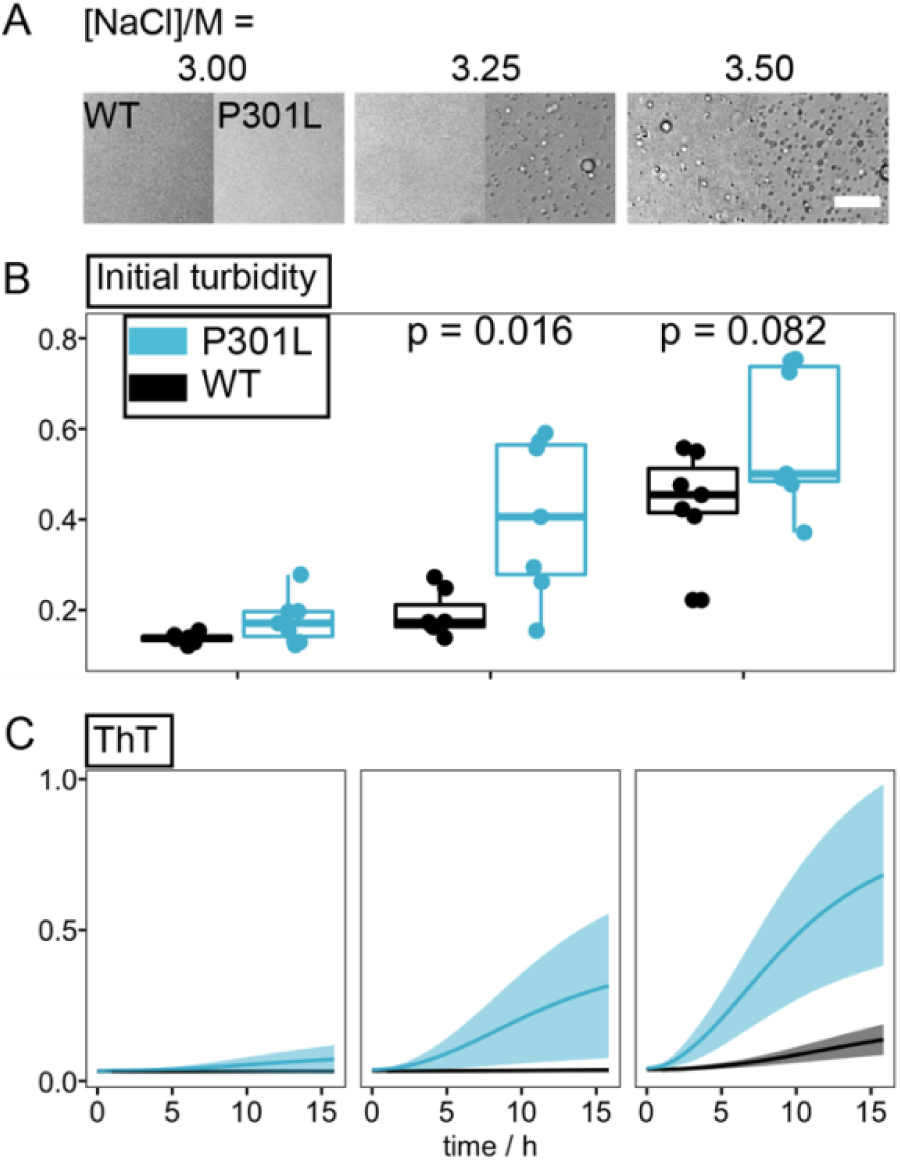
Effects of P301L mutation on LLPS and amyloid aggregation of tau at high salt concentration. **A**. Representative microscope images of tau187 Cysless and CyslessP301L mutants at varying [NaCl]. **B**. Turbidity of corresponding samples in A. **C**. ThT fluorescence of samples in A after overnight incubation at room temperature. ThT fluorescence was scaled by a constant of 2×10^4^ throughout the manuscript. Ribbons in C show standard deviation of readings from 5 different batches of proteins.

### Dehydration facilitates hydrophobically driven LLPS-HS

To reveal the mechanisms that promote the transition of tau from high salt droplet to tau in fibrils, we next focused on investigating the driving forces for tau to form in LLPS-HS. The dissolution by 1,6-HD showed high salt droplets to be held together by hydrophobic interaction (**Figure 1**D). When increasing the amount of 1,6-HD, we found the minimum [NaCl] to induce LLPS-HS to shift significantly towards higher NaCl concentration (**Figure 4-figure supplement 3**C). This implies that a greater amount of salt is needed to restore the same enhancement of hydrophobic interactions that are disrupted by 1,6-HD.

Turbidity shows LLPS-HS is favored at higher temperature, following lower critical saturation temperature behavior (LCST) (**Figure 2-figure supplement 1**A, C). This suggests that free energy for LLPS-HS is rendered negative (ΔG<0) by a dominant and positive entropy term, TΔS, according to ΔG = ΔH-TΔS. LCST behavior is seen when phase separation is accompanied by an increase in total entropy (i.e. ΔS>0) that, as a result, leads to an increasingly dominant and positive entropy (TΔS) term at higher temperature. The temperature at which ΔG = 0 is then the phase transition temperature, cloud point T_cp_, above which phase separation occurs spontaneously. Interestingly, either increasing [NaCl] or increasing [tau] lowers the T_cp_ above which LLPS-HS occurs (**Figure 4-figure supplement 1**). This finding suggests that both the addition of salt and higher tau concentration contribute to entropy gain upon LLPS. When adding PEG as a molecular crowder, we observed that the minimum [NaCl] necessary to induce LLPS-HS also shifted towards lower concentrations (**Figure 4-figure supplement 3**B). A recent study by Park et al. demonstrated that PEG increases the entropy term upon LLPS by dehydration, and so enhances LLPS without partitioning into the dense phase of LLPS [30]. Taken together, our results are consistent with NaCl and PEG both sequestering water, and hence depleting the protein system of free water in a process known as salting out [41], [59]. Dehydration of the tau-water system will facilitate hydrophobic interactions and subsequent condensation by an increase of the entropic term TΔS.

While the observations made by systematic changes in [NaCl], [tau], [PEG] and temperature is consistent with dehydration- and entropy-driven LLPS-HS, a more direct observation of dehydration is more satisfying. To measure the effect of dehydration near the tau protein surface, we labeled tau with unpaired electron spin labels to perform electron spin envelope echo modulation (ESEEM) and Overhauser dynamic nuclear polarization (ODNP) measurements. ESEEM measures the relative changes in the local water density within a ∼3-6 Å shell around a spin label [60], [61], while ODNP measures water accessibility [62] and translational diffusivity within a shell of ∼5-10 Å [63], [64] (**Figure 4**A).

We labeled tau187 with MTSL on site 313 within the R3 domain and dissolved it into a deuterium-based buffer. The magnitude of the ESEEM deuterium peak intensity dropped by over 20%, immediately upon addition of NaCl that led to the formation of high salt droplets (**Figure 4**B). This result represents a significant reduction of local water density near site 313.

We furthermore carried out ODNP relaxometry measurements to determine the cross-relaxivity parameter k_σ_, for tau187 at sites 303, 313, 322 and 404, representing the 4R domain and the C-terminal region of tau. The k_σ_ value reflects on local water accessibility and/or translational diffusivity within ∼1 nm of the spin label (Eq. 5 of [65]). We found that the value of k_σ_ dropped from 31 - 56 M^-1^s^-1^ to 9 - 23 M^-1^s^-1^ for all four sites as soon as high salt droplets formed (see per site trend in **Figure 4**C). Note that k_σ_ measured on the free radical 4-hydroxyl-TEMPO was unchanged upon addition of salt at concentrations that induce LLPS-HS (**Figure 4**C). Therefore, the decrease of k_σ_ observed reflects a change in hydration properties near the tau protein surface upon LLPS-HS formation. The reduction of the deuterium peak magnitude in ESEEM at site 313 shows that the reduction of water accessibility contributes to the drop of k_σ_. Given the ∼25% reduction of deuterium peak magnitude together with the 50∼80% drop in k_σ_ (**Figure 4, Figure 4-source data 1**), it is reasonable to assume that it reflects, in addition to local dehydration, a significant slowdown of surface water diffusivity near the microtubule binding domain and C-terminal of tau when condensed into LLPS-HS. Strikingly, k_σ_ remains unchanged upon formation of tau-RNA LLPS-ED (**Figure 4**C), showing that dehydration of LLPS-internal water is a characteristic associated with hydrophobically driven LLPS-HS, but not of LLPS-ED.

### Dehydration reduces tau dynamics and triggers oligomer formation

To study the dynamics of tau within high salt droplets, we carried out continuous wave (cw) EPR spectroscopy. We labeled tau187 at four different sites: 303, 313, 322 and 404, and extracted the rotational correlation times of the tethered spin label, t_R_, from fitting the cwEPR lineshape (see Materials and Methods). The value for t_R_ of the tethered spin reflects on the local protein mobility, as the spin label is tethered to the protein side chain with steric constraints [66]. We first looked at t_R_ of the four sites before and after forming LLPS-HS. We found that LLPS-HS formation results in dramatic changes in the spectral lineshape of the spin labeled tau (**Figure 5**A, **Figure 5-figure supplement 2**). We used the microscopic order macroscopic disorder (MOMD) model [67] with incremental complexity to fit the cw EPR lineshapes, using the MultiComponent program developed by Christian Altenbach. At [NaCl] = 0–1.5 M (no droplets), lineshapes of the samples were well fit with one isotropic t_R_ component (**Figure 5-figure supplement 2**). Increasing [NaCl] to 3 M and above resulted in LLPS-HS. The lineshape of the samples deviated from the one component fit, and required an additional mobility component (**Figure 5-figure supplement 1**A). The extracted t_R_ and the population of each component is represented in **Figure 5**B as a function of salt concentration. We found that t_R_ of the second component was significantly higher than for the first component (i.e. slower dynamics), for all sites. The second component represented 65-96% of the spin label population above 3 M NaCl (**Figure 5-source data 1**). Meanwhile, increasing [NaCl] up to 4.5 M alone does not change the lineshape of the free radical 4-hydroxyl-TEMPO (**Figure 5-figure supplement 4**), showing the effects of high salt concentration do not change the rotational dynamics of the spin itself. We hypnotize that this increase of rotational correlation time t_R_ reflect on the formation of higher molecular weight assemblies, i.e. oligomers. These results imply that the dynamics of the majority of tau proteins, reported on via protein-tethered spin labels, is slowed down in the LLPS-HS state. In addition to the slowed rotational dynamics, we found emergence of rotational anisotropy in the second component for [NaCl] above 4 M. The presence of anisotropy indicates that the tethered spin label is highly restricted [67], [68], likely from steric hindrance of nearby tau protein, suggesting that tau starts to more tightly pack into protein assembly at very high salt concentration.

Next, we endeavor to show whether the formation of these oligomer species, revealed by slow label dynamics, originates from high salt concentration or from LLPS itself. First, in the absence of high salt droplets, we observed that the lineshape of cw EPR spectra of the tau-tethered spin label broadened upon NaCl addition, giving an average t_R_ of ∼450 ps without NaCl and ∼600 ps at 1.5M, at all sites (**Figure 5-source data 1**). In addition, at constant [NaCl] we titrated [HD] from 0% to 10%, where [HD] above 2% eliminated droplets. CwEPR lineshape analysis showed that the correlation time was neither abruptly nor distinctly reduced in the absence of droplets above 2% HD (**Figure 5-figure supplement 3**). These results show that the drastic reduction in protein dynamics, which we hypothesized to reflect on the formation of oligomers, originate from dehydration by concentrated salt, only indirectly by LLPS.

### P301L promotes hydrophobic interactions

One of the mysteries in tau amyloid aggregation is how a single site mutation, e.g, P301L, dramatically promotes the aggregation propensity of a protein of hundreds of amino acids. We propose here a plausible mechanism. We utilize LLPS-HS as a tool to measure hydrophobic interactions among tau. We studied the effects of the P301L mutation on LLPS-HS propensity. We measured turbidity of tau187 with and without P301L mutation under the exact same conditions. Turbidity from independent biological repeats showed that in the presence of an intermediate salt concentration of 3.25 M, the presence of P301L mutation significantly promoted LLPS-HS formation (**Figure 6**B). In other words, the P301L mutation lowers the [NaCl] threshold required to induce LLPS-HS. Representative microscope images, shown in **Figure 6**A, confirmed that P301L promotes high salt droplet formation. These results suggest that the P301L mutation promotes hydrophobic interactions among tau proteins, in agreement with previous proposal that P301L mutation extends local conformation, hence reveals the hydrophobic hexapeptide _306_VQIVYK_311_ [69]. However, whether the increased hydrophobicity of tau in the presence of P301L is solely due to conformational changes or other biophysical factors requires in-depth studies in the future. In addition, we observed that the ThT activity triggered by LLPS-HS (shown in **Figure 3** for wild type (WT)) increases further for P301L compared to WT (**Figure 6**C). This enhanced ThT fluorescence for P301L mutant is most likely a combination of both the promotion of droplet formation (which in turn triggers ThT-sensitive aggregation) and a high aggregation propensity that facilitate the droplet to fibril transition.

## Discussion

Protein liquid-liquid phase separation in pathological contexts have led to intense investigation in the recent past. In particular, tau LLPS has been studied in several reports leading to contrasting hypothesis whether or not LLPS directly triggers amyloid aggregation. We address in this study two key questions: (1) Is tau LLPS of a universal type or could droplets formed in different conditions possess different properties? (2) Is there a direct cause-and-effect relationship between LLPS and aggregation?

In this report, we have shown that tau at high salt concentration is capable of forming LLPS (referred to as LLPS-HS) driven by hydrophobic interactions. High NaCl concentrations both screen electrostatic interactions and sequester water away from the protein, thereby amplify the effect of hydrophobic association of tau. The concentration of salt needed to achieve LLPS of tau of the order of ∼3M is not physiological by any means. However, the influence of such condition on the proteins or biopolymers, i.e. strong electrostatic screening and amplified hydrophobic attraction can easily represent physiologically relevant state. For example, tau under conditions of significant phosphorylation and acetylation would experience effects similar to that of screening the net positive charge effects of the many lysine residues lining on the tau surface, while strong hydrophobic attraction was proposed to be at the origin of hyperphosphorylated tau aggregation [70].

Using EPR spectroscopy, FILM and biochemical assays, we showed that LLPS-HS exhibit slowed local dynamics, restricted water accessibility, elevated microviscosity, as well as irreversible maturation, in stark contrast to electrostatically driven LLPS-ED. Many different conditions can facilitate tau LLPS, but this study shows that the classification of hydrophobically or electrostatically driven LLPS might be an effective framework to classify tau droplets and predict their properties or effects on aggregation. We used tau-RNA complex coacervation to showcase LLPS-ED, but as shown in **Figure 1**D and elsewhere [33], [34], both complex and simple coacervation can form LLPS-ED.

Establishing a causal relationship between LLPS and protein aggregation is challenging because the two processes can occur under identical conditions without necessarily being dependent [38]. Importantly, under conditions where both phenomena occur, it is tempting to deduce a cause-and-effect relationship. Firstly, droplets can form and dissolve within seconds to minutes, while aggregation occurs on the timescale of hours to days, therefore the sequential occurrence of droplets then aggregates does not necessarily imply LLPS is an intermediate state towards aggregation. Secondly, droplets, by definition, increase protein concentration. Therefore, even if both phenomena were completely independent, one would observe aggregates slowly forming in the inside of pre-existing droplets, merely because it is where the protein is, which might lead to the wrong conclusion that droplets directly *trigger* aggregation. In our opinion, a robust way to establish causation between droplets and aggregates is to examine multiple conditions across the LLPS phase boundary, and correlate the changes of droplet volume to the change of fibril mass. Using this approach, we have shown in [38] that although tau-cofactor LLPS-ED and amyloid aggregation are concomitant in many conditions, they are biophysically independent processes. In this study, we show that hydrophobically driven LLPS-HS is different, and that LLPS-HS of tau trigger amyloid formation (**Figure 3**C and **Figure 3-figure supplement 3**).

Previous reports on tau LLPS made with K18 at low salt concentrations showed that the droplets were dissolved above ∼200 mM NaCl and at 3 % HD, suggesting that a combination of electrostatic and hydrophobic driving forces were holding the liquid condensed state together [19], [51]. These droplets were not reported to evolve spontaneously to ThT-active species. In contrast, droplets formed by hyperphosphorylated 2N4R in Wegmann *et al*. [20] were shown to evolve to amyloid fibrils and to be insensitive to [NaCl]>1M, but to be sensitive to 10% HD, showing that they are driven mostly by hydrophobic interactions. These results together with our work suggest a general mechanism that hydrophobically-driven droplets promote aggregation.

We have shown that the local environment in the high salt droplets is drastically different from the one in tau LLPS-ED droplets. FLIM showed that, initially after high salt droplet formation, i.e. even in the absence of significant aging, the micro-viscosity is significantly higher in LLPS-HS than in LLPS-ED droplets. Furthermore, ESEEM and ODNP measurements revealed that water accessibility and water dynamics is significantly more retarded in LLPS-HS than in LLPS-ED. These properties found in LLPS-HS are favorable to protein aggregation because amyloid formation requires a critical step of dehydration, where even hydrophilic residue must give up their ideal hydration to pack into cross-β sheets and steric zippers. Besides, amyloid fibers form a solid-like matrix of high viscosity. We suggest that the features of high viscosity, perturbed protein dynamics and dehydration are general hallmarks of protein LLPS on-pathway to aggregation.

Cw EPR linseshape analysis provides a mechanistic view of tau aggregation through LLPS-HS. We found that increasing the concentration of salt progressively triggers the formation of high order oligomers, as evidenced by a long rotational correlation time of the spin labels tethered to tau. These oligomers are observed even at medium [NaCl] (e.g. 1.5 M) where droplets are not present. An analysis of the dipolar broadening of the cwEPR spectra (**Figure 5-figure supplement 4**) shows the absence of specific packing (such as in-register cross-β sheets or steric zipper structures) in these ThT-inactive oligomers. LLPS seems however necessary and sufficient to convert these oligomers into ThT-active species and eventually amyloid fibers, as shown by the direct correlation between ThT fluorescence and droplet amount (**Figure 3**C).

Finally, we demonstrate that the disease-associated mutation P301L promotes both LLPS-HS and amyloid formation (**Figure 6**). Conversely, P301L mutation has no influence on the formation of LLPS-ED [38]. Therefore, we can conclude that P301L mutation promotes intermolecular hydrophobic interactions that drives LLPS-HS. Furthermore, the correlation plots shown in **Figure 3**C and **Figure 6-figure supplement 2**C show that at similar droplet quantity, P301L mutant will form greater quantity of ThT active species, and does so more rapidly than tau WT. The aggregation-promoting effect of P301L is well known and elegant explanations have been suggested recently [69, p. 201]. Here, we show that the P301L mutation exerts two seemingly independent effects that both ultimately increase aggregation: (i) It promotes local hydrophobic interactions, thereby promoting hydrophobically-driven phase separation, which in turn triggers aggregation and (ii) promotes aggregation-prone conformations, in particular by exposing amyloidogenic segments PHF6(*), independently of LLPS formation. Other disease-related mutations might not combine these effects, because they do not modulate the accessibility of the hydrophobic PHF6(*) segments, explaining why P301L is particularly potent at promoting fibril formation under wide experimental conditions, from *in vitro*, in cells and *in vivo* mouse models.

In conclusion, this study uncovers that there is a clear cause-effect relationship between LLPS and aggregation of tau, only when LLPS is driven by hydrophobic association of tau that we found to be preceded by dehydration of its interfacial water. Different physical or biological factors can drive hydrophobically driven LLPS. While this study showcases the principle by relying on dehydration-promoting salting-out mechanisms induced at high [NaCl] concentration, we posit that hydrophobically driven LLPS can be exerted by many other factors of pathological significance, including the posttranslational modification of tau that can mute the strong positive surface charge density from the abundant lysine residues on tau, e.g. by acetylation and phosphorylation, or enhanced intracellular osmotic crowding pressure, as well as certain pathological mutations of tau, with P301L representing one prominent example. Hence, the liquid condensed state of tau that can be seen *in vitro* and intra cellularly may be a mediating mechanism by which tau can be sequestered and initially protected from aggregation if LLPS can be reversed, or is inadvertently and irreversibly on pathway towards a pathological state.

## Acknowledgements

The authors acknowledge support for the studies of LLPS mechanisms from the National Institutes of Health (NIH) (grant R01AG056058 (S.H.) and support of the Tau consortium of the Rainwater foundation for SH and KSK to study the effect of LLPS on tau pathology. We acknowledge support for the synthesis of BODIPY-NHS dye from the National Science Foundation (DMR-1922042). We acknowledge the use of the NRI-MCDB Microscopy Facility and the Resonant Scanning Confocal supported by NSF MRI grant DBI-1625770 at UC, Santa Barbara. We acknowledge the use of the MRL Shared Experimental Facilities which are supported by the MRSEC Program of the NSF under Award No. DMR 1720256; a member of the NSF-funded Materials Research Facilities Network (www.mrfn.org)

## Materials and Methods

### Tau proteins expression and purification

Both full length human tau (2N4R) and its N-terminal truncated, microtubule binding domain contained variant tau187 (residues 255-441 with a His-tag at the N-terminus) were used for *in vitro* studies. The cloning, expression, and purification of both 2N4R and tau187 have been previously described [71], [72]. The cysteine-free variants of both 2N4R and tau187 (C291SC322S) were generated via site-direct mutagenesis.

E. coli BL21 (DE3) cells previously transfected were cultured from frozen glycerol stock overnight in 10 mL luria broth (LB) which was used to inoculate 1 L of fresh LB. Culturing and inoculation were performed at 37 °C with shaking of 200 rpm. At OD_600_ of 0.6–0.8. After cell growth, 2N4R culture was incubated with 500 mM NaCl and 10 mM betaine before inducing expression. Both 2N4R and Tau187 variant expressions were induced by incubation with 1 mM isopropylß-D-thiogalactoside (Sigma Aldrich) for 2–3 h. Cells were harvested by centrifugation for 10 min at 6000 × g (Beckman J-10; Beckman Instruments, Inc.), and the pellets were stored at −20 °C until further use.

Cell pellets were resuspended in lysis buffer (Tris-HCl pH 7.4, 100 mM NaCl, 0.5 mM DTT, 0.1 mM EDTA, 1mM PMSF) with 1 Pierce protease inhibitor tablet (Thermo Fisher, EDTA-Free). Lysis was initiated by the addition of lysozyme (2 mg/ml), DNase (20 µg/ml), and MgCl_2_ (10 mM) and incubated on an orbital shaker for 30 min in room temperature. Lysate was then heated to 65 °C for 13 min, cooled on ice for 20 min and then centrifuged to remove the precipitant. The supernatant was loaded onto a Ni-NTA agarose column pre-equilibrated with wash buffer A (20 mM sodium phosphate pH 7.0, 500 mM NaCl, 10 mM imidazole, 100 µM EDTA). The column was then washed with 20 ml of buffer A, 15 ml buffer B (20 mM sodium phosphate pH 7.0, 1 M NaCl, 20 mM imidazole, 0.5 mM DTT, 100 µM EDTA). Purified tau was eluted with buffer C (20 mM sodium phosphate pH 7.0, 0.5 mM DTT, 100 mM NaCl) supplemented with varying amounts of imidazole increasing from 100 mM to 300 mM. The protein was then concentrated via centrifugal filters (MWCO 10 kDa; Millipore Sigma) and the buffer was exchanged into final buffer by PD-10 desalting column (GE Healthcare). The final protein concentration was determined by UV-Vis absorption at 274 nm using an extinction coefficient of 2.8 cm^-1^mM^-1^ for tau187 and 5.8 cm^-1^mM^-1^ for 2N4R, calculated from absorption of Tyrosine [3]. Same purification protocols were used for both 2N4R and tau187.

### Bright field microscopy and TEM imaging

To confirm the occurrence of LLPS and the formation of liquid droplets, bright field microscope images were taken immediately after sample preparation, using an inverted compound microscope (Zeiss Axtrovert 200M).

For TEM imaging 10 μL of recombinant tau fibril samples were applied to copper grid (Electron Microscopy Science, FCF-300-Cu) cleaned with plasma for 20 s. Samples were stained with 10 μL 1.5 w/v % uranyl acetate. Grids were then imaged with a JEOL JEM-1230 (JEOL USA, Inc) at the indicated magnifications.

### Turbidity measurements

Turbidity of samples at varying [NaCl] were represented by OD_500_ measured using a plate reader (Bio-Tek Synergy 2) and 384-well microplate (Corning 3844). To minimize the inference of ThT fluorescence and tau fibril scattering, averaged absorbance of the initial 10 min where ThT fluorescence is below 10% was used as turbidity reading.

Turbidity of samples at ramping temperatures were represented by OD_500_ measured using Jasco J-1500 CD Spectrometer (JASCO Inc.) equipped with temperature controller and spectrophotometer. 120 μL of tau in designated conditions were prepared in a 100 μL cuvette (Starna Scientific Ltd) and kept at 4 °C for 5 min before cycling. Heating and cooling temperatures were ramped at 1 °C/min while OD_500_ was monitored.

### ThT assay and half time analysis

LLPS samples were prepared onto a 384-well microplate with additional 1 µM Thioflavin T. ThT fluorescence was read from a plate reader (Bio-Tek Synergy 2, excitation 440/30, emission 485/20, number of flash 16). Data was processed using R package gen5helper following the previous reported methods to reduce artifacts caused by the lamp fluctuation [38].

We followed the previously reported methods [38] to extract the half time and maximum ThT fluorescence, by scaling ThT fluorescence to 0 to 1 and fitting to a sigmoid function,

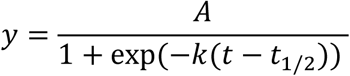

where the fitted A and t_1/2_ were used as maximum ThT fluorescence and half time, respectively. Fitting results were filtered by p < 0.05 to ensure a good fit.

### Synthesis of BODIPY-NHS Dye

BODIPY-NHS was synthesized as previously described with few modifications [55]. A brief description for the synthesis of BODIPY-NHS dye is provided in **Figure 2-figure supplement 3** and full details can be found at DOI:<u>10.6084/m9.figshare.12746756</u>. BODIPY-NHS can be synthesized in six steps starting from 4-hydroxybenzaldehyde. In the first step, etherification of 4-hydroxybenzaldehyde by potassium carbonate and ethyl 4-bromobutyrate afforded ethyl 4-(4-formylphenoxy)butyrate **(1)** in 98% yield. Then, hydrolysis with potassium hydroxide provided 4-4(formyl-phenoxy)butyric acid **(2)** in 96% yield. In order to overcome solubility issues that the carboxylic acid intermediate presents in pyrrole, used in step 4, it was converted to the corresponding silyl ester. It reacted with tert-butyl(chloro)diphenylsilane in a reaction promoted by triethyl amine and 4-dimethylaminopyridine affording tert-butyldiphenylsilyl 4-(4-formylphenoxy)butanoate **(3)** in 68% yield. The aldehyde **(3)** was then treated with TFA (12 mol%) and an excess of pyrrole under solvent-free conditions providing the dipyrromethane **(4)** in 66% yield. Oxidation with 2,3-dichloro-5,6-dicyano-1,4-benzoquinone followed by reaction with BF_3_·OEt_2_ and triethylamine lead to BODIPY-COOH in 36% yield. Steglich esterification with N-hydroxysuccinimide yielded BODIPY-NHS (64% yield).

### Conjugation of BODIPY-NHS to 2N4R

2N4R protein was diluted to a concentration of 20 µM in a PBS buffer. 20 mM BODIPY dissolved in DMSO was added to the protein to a final concentration of 20 µM. The solution was stirred at room temperature for 4 hours and any unreacted BODIPY-NHS was quenched by adding a solution of excess glycine. The unreacted dye was separated from the conjugated protein with a 7k MWCO Zeba™ Spin Desalting Column equilibrated with a 20 mM HEPES buffer, pH = 7.0. The conjugated protein was concentrated in an 3K MWCO Amicon Ultra Spin Column to a final concentration of approximately 500 µM. Protein concentration was determined with a Pierce Assay, while the amount of conjugated dye present was calculated by measuring its absorbance at 510 nM. Typical labeling reactions gave a 1:1 labeling efficiency.

### Measurement and Analysis of FLIM Spectra

All FLIM measurements were made on a Leica SP8 FALCON Resonant Scanning Confocal Microscope equipped with TCSPC. Images were recorded such that each pixel had a minimum of 1000 photons collected. All data analysis was performed with FLIMFIT [73]. As observed previously [58], lifetime decay curves required multiexponential fits. We found that the higher lifetime component of a 2-exponential fit tracked well with control viscosity measurements made in varying glycerol concentrations and as such it was used for all subsequent analysis.

### CwEPR and lineshape analysis

In this report, we used four variants of tau187 for cwEPR, ODNP and ESEEM studies: tau187/C291S/C322S/S303C, tau187/C291S/C322S/G313C, tau187/C291S, tau187/C291S/C322S/S404C, with one cysteine at site 303, 313, 322 and 404, respectively.

We followed the previous reported methods [38] for spin labeling, cwEPR acquisition, and lineshape analysis. For spin labeling, the freshly eluted variants above were replaced in 20 mM HEPES pH 7.0, using a PD-10 desalting column (GE Healthcare). Protein after PD-10 was labeled overnight at 4°C by immediately mixing with a 10-fold molar excess of the spin label (1-oxyl-2,2,5,5-tetramethylpyrroline-3-methyl) methanethiosulfonate (MTSL; Toronto Research Chemicals), resulting in spin labelled tau at designated site. Excess label was removed using PD-10. The protein was concentrated using centrifugal filter (MWCO 10 kDa; Amicon) and the final protein concentration was determined by UV-Vis absorption at 274 nm as mentioned above. Cystein-free non-labeled tau187C291SC322S was used in order to achieve spin dilution.

Cw EPR measurements were carried out using a X-band spectrometer operating at 9.8 GHz (EMX; Bruker Biospin, Billerica, MA) and a dielectric cavity (ER 4123D; Bruker Biospin, Billerica, MA). Spin labeled tau187 was mixed with tau187C291SC322S at 1:4 molar ratio to reach 20% spin labeling. A sample of 4.0 μl volume was loaded into a quartz capillary (CV6084; VitroCom) and sealed at both ends with critoseal, and then placed in the dielectric cavity for measurements. Cw EPR spectra were acquired by using 1.8 mW of microwave power, 0.5 gauss modulation amplitude, 100 gauss sweep width, and 8-64 scans for signal averaging.

The recorded cw EPR spectra were fit with simulation using models with incremental complexity. EPR simulation and fitting were performed using MultiComponent, a program developed by Christian Altenbach (University of California, Los Angeles). For all spectra fitting, the magnetic tensors A and g were fixed and used as constraints as previously reported [74]. These values are A_xx_ = 6.2 G, A_yy_ = 5.9 G, A_zz_ = 37.0 G, and g_xx_ = 2.0078, g_yy_ = 2.0058, and g_zz_ = 2.0022. For soluble tau and tau at [NaCl] < 2.0 M, the cw EPR spectra of all four spin-labeled tau variants were best fitted with a single isotropic component simulation and the rotational diffusion constant (R) can be extracted. The rotation correlation time τ_R_ was calculated using τ_R_ = 1/(6R). For tau at 2.0 M < [NaCl] < 4.0 M, the single isotropic component simulation failed to capture the lineshape (**Figure 5-figure supplement 1**A). A second isotropic component was added to increase the complexity of the model. The second isotropic component were set to be identical to the first component, except with an independent R value. The fitting parameters were limited at a minimum, which includes the population, p and the rotational diffusion constants of the first and second isotropic component R_1_ and R_2_. For tau at [NaCl] > 4.0 M, both the signal and double isotropic component simulation failed to capture the lineshape (**Figure 5-figure supplement 1**B). Anisotropic parameters were added to the second components and subjected to fit. The fitting parameters include p, R_1_, R_2_, and the added parameters for the second components including the tilt angle of the diffusion tensor α_D_, β_D_, orienting potentials c_20_, c_22_, c_40_, c_44_. Fitted spectra and parameters of all four sites at different [NaCl] are shown in **Figure 5-figure supplement 2** and **Figure 5-source data 1**, respectively.

### Overhauser dynamic nuclear polarization (ODNP)

We used the aforementioned four spin labeled tau187 variants (singly labeled on site 303, 313, 322 and 404, respectively), and followed previously reported methods for ODNP experiments [30]. 100 µM spin labeled tau187 was mixed with designated concentration of NaCl. Immediately after sample preparation, 3.5 µL of the samples were loaded into quartz capillaries of 0.6 mm ID × 0.84 mm OD (Vitrocom, New Jersey, USA), and two ends of the tubes were sealed with Critoseal and beeswax respectively. ODNP experiments were performed using a Bruker EMXPlus spectrometer and a Bruker Avance III NMR console (Bruker, Massachusetts, USA). The capillary tube was mounted on a home-built NMR probe with a U-shaped NMR coil, and was set in a Bruker ER 4119HS-LC sensitivity cavity. Samples were irradiated at 9.8 GHz with the center field set at 3484 G and sweep width of 120 G. Dry air was kept at temperature of 18 °C and purged through the NMR probe during all measurements. ODNP data was analyzed using customized python script WorkupODNP and ODNPLab, available on https://www.github.com. Theory of ODNP and details in the experiment are previously reported in other studies [63], [64].

### 3-Pulse Electron Spin Echo Envelope Modulation (ESEEM)

We followed previously reported methods for ESEEM measurements [75]. ESEEM measurements were performed at X-Band on an ELEXSYS E580 spectrometer at 80 K with an Oxford instruments cryostat (CF935P) and temperature controller (ITC503). A Bruker MS3 resonator was maximally overcoupled to a Q-factor of approximately 100 for these measurements. Immediately after sample preparation, 35 μL of sample was loaded into a 3 mm OD quartz EPR tube and flash frozen in liquid nitrogen before being inserted into the resonator. Spin labeled tau187 at site 313 was used for measurement. All experiments were carried out in deuterated 20 mM HEPES with 20 wt% Ficoll.

The ESEEM data was fit by 5^th^ order polynomial to correct the background decay, hamming windowed, zero-filled and Fourier-transformed, following reported method [75]. We interpreted the reduce of the the intensity of the deuterium peak as a result of dehydration near the site 313. Raw data was shown in **Figure 4-figure supplement 2**.

### Estimate Unbound Tau Concentration

Assuming tau binding to microtubule has a stoichiometry of 1:1 between tau molecule and tubulin monomer, regardless of different types of tubulin monomers, the dissociation process can be expressed as

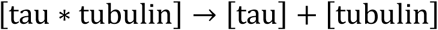

where [tau*tubulin] and [tau] represent the bounded and unbounded tau concentration, respectively. Knowing

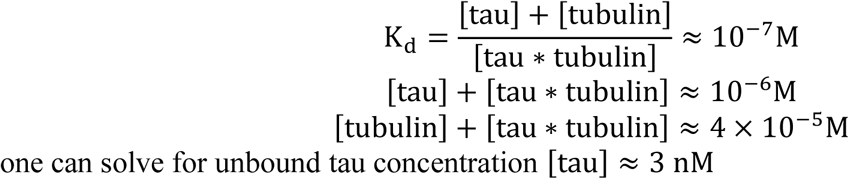

one can solve for unbound tau concentration [tau] ≈ 3 nM

## Figure and Table Captions

**Figure 1-figure supplement 1.**
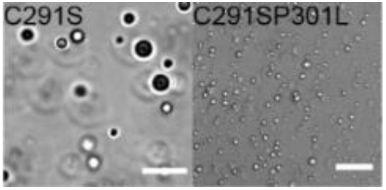
Microscope images of LLPS-high salt of various tau constructs. 450 µM Tau187C291S with 2.2 M NaCl; 100 µM tau187C291SP301L with 3.0 M NaCl.

**Figure 2-figure supplement 1.**
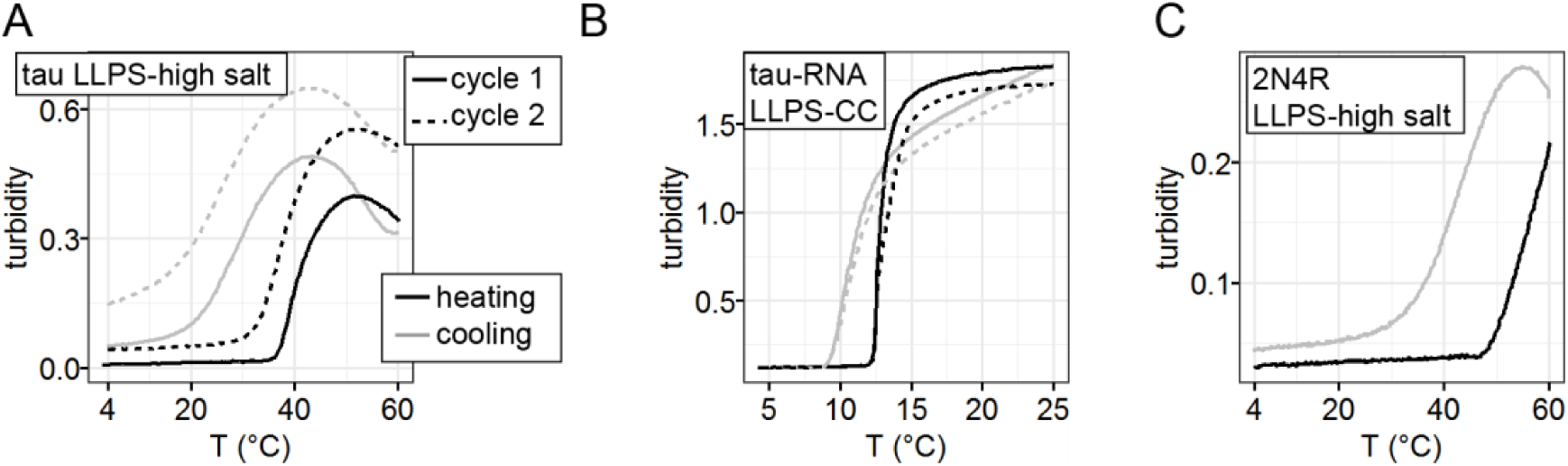
Temperature dependence of LLPS-HS. **A**. turbidity of tau187 LLPS-high salt upon heating-cooling cycles. 34 μM tau187 and 3.5M NaCl was used. **B**. turbidity of tau187-RNA LLPS-CC upon heating-cooling cycles. 100 μM tau187 with 300 μg/mL PolyU RNA and 30 mM NaCl was used. **C**. turbidity of 2N4R LLPS-high salt upon heating-cooling cycle. 20 μM tau with 3.5M NaCl was used. Samples were pre-cooled to 4 °C, heated to 25 or 60 °C then cooled back to 4 °C (cycle 1, solid lines). A second cycle immediately follows (cycle 2, dashed lines). The rate of both heating and cooling are 4 °C/min in A and C, and 1 °C/min at B.

**Figure 2-figure supplement 2.**
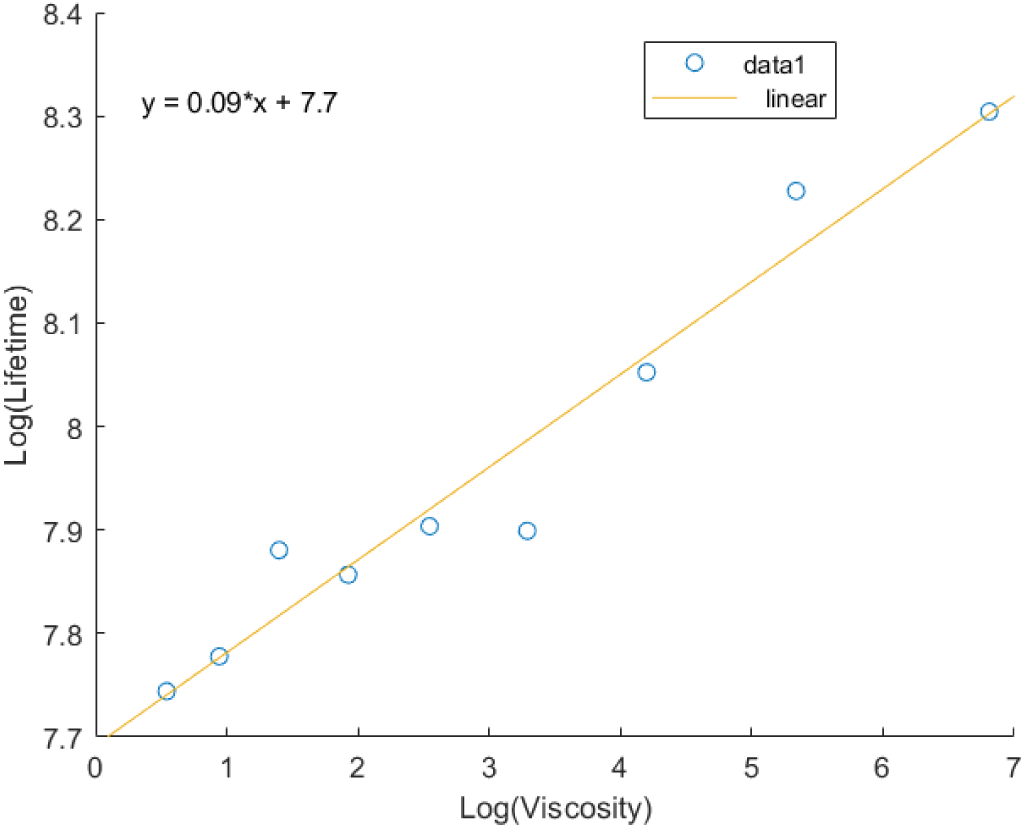
Calibration and Fit of Fluorescent Lifetimes to a Glycerol Standard Curve. 4 µM 2N4R conjugated the BODIPY was placed in solutions of glycerol concentrations in ddH_2_O ranging from 20-100%. FLIM measurements of the solutions were made and the resulting gaussians were fit and extracted in FLIMFit. The gaussians were fit and the mean values were plotted on a log/log scale against the corresponding solutions viscosity. The data was fit to the Förster Hoffman equation,

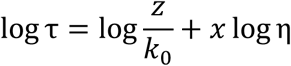

Where τ is the lifetime in ps, 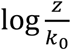 is treated as a variable, x is the slope, and η is the viscosity in centipoise. Fitting the data gave an equation of the form *y* = 0.09*x* + 7.7.

**Figure 2-figure supplement 3.**
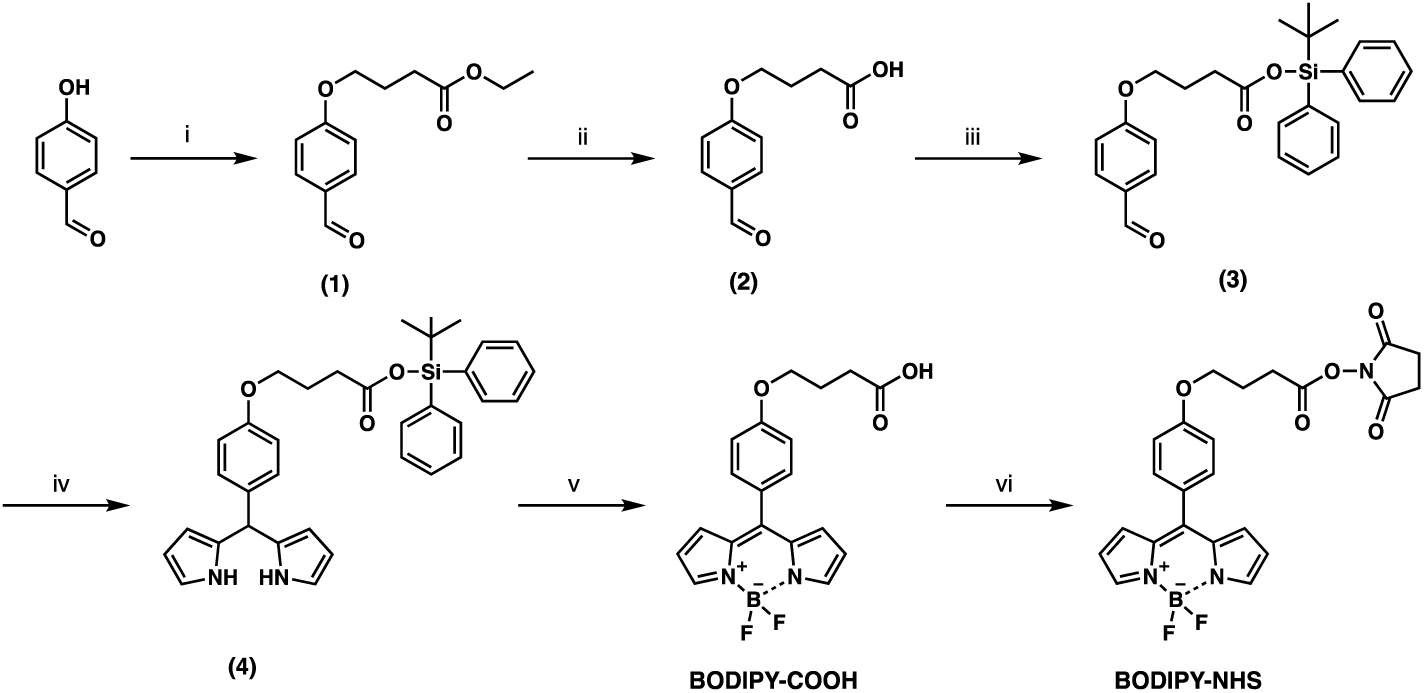
Synthesis scheme of BODIPY-NHS: (i) ethyl 4-bromobutyrate, K_2_CO_3_, DMF, rt, 24 h, 98% yield; (ii) 2N KOH, MeOH, reflux, 2 h, 96% yield; (iii) DMAP, Et_3_N, THF then tert-butyl(chloro)diphenylsilane, 0°C to rt, 20 h, 68% yield; (iv) pyrrole, TFA, rt, 1 h, 66% yield. (v) DDQ, CH_2_Cl_2_, rt, 1 h and then BF_3_·(OEt_2_), Et_3_N, rt, overnight, 36% yield; (vi) N-hydroxysuccinimide, DCC, DMAP, THF, 0°C to rt, overnight, 64% yield.

**Figure 3-figure supplement 1.**
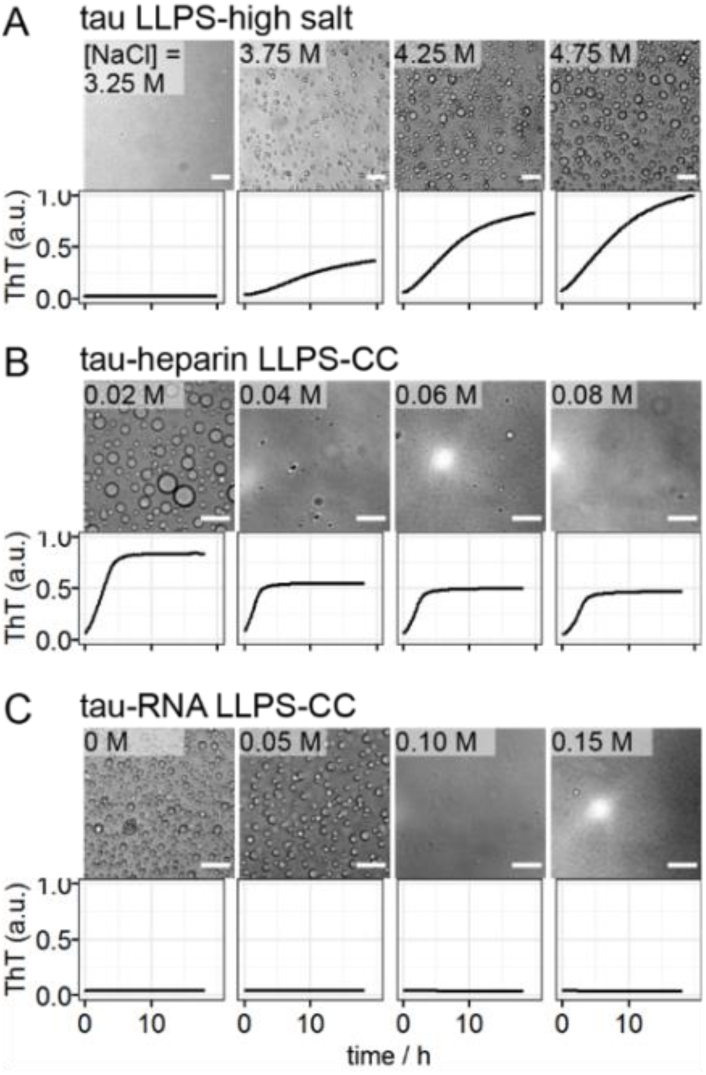
Representative microscope images and ThT fluorescence at room temperature overnight of tau LLPS-high salt, tau-heparin LLPS-CC, tau-RNA LLPS-CC at various [NaCl]. 44 µM tau187, 73 µg/mL heparin and 132 µg/mL RNA were used.

**Figure 3-figure supplement 2.**
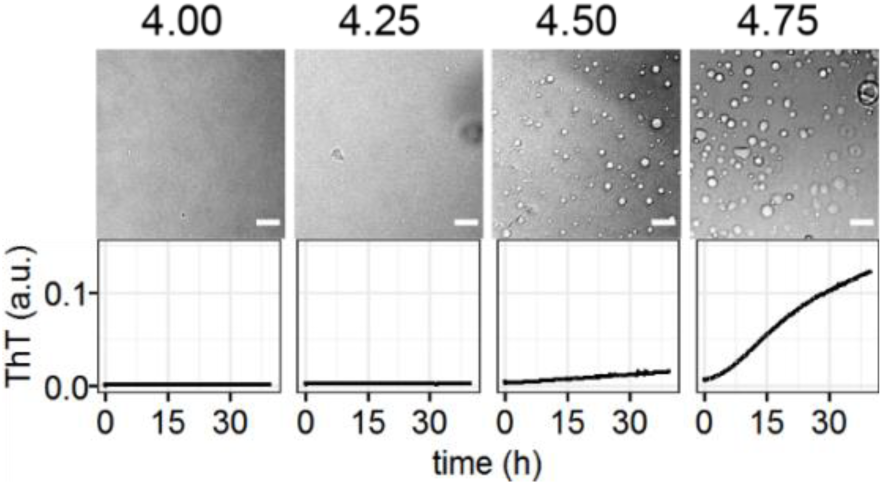
Correlation of LLPS-high salt and amyloid aggregation of 2N4R. 20 μM 2N4R was incubated with varying [NaCl]. Microscope images were taken 10 minutes after mixing while ThT fluorescence readings were recorded overnight. Scale bar length was 25 μm.

**Figure 3-figure supplement 3.**
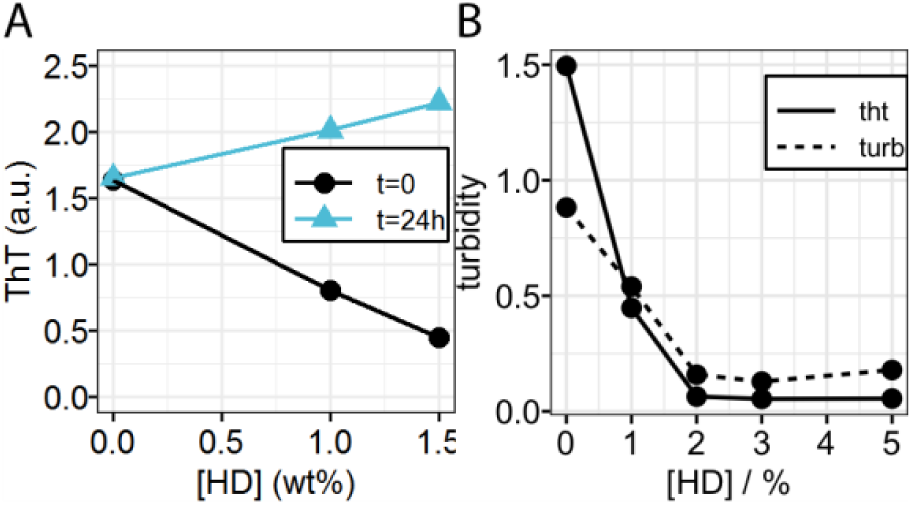
Hexanediol dependence of ThT fluorescence and turbidity of LLPS-HS. **A**. Varying concentration of 1,6-hexanediol were added to fresh LLPS-HS sample (t=0) and matured LLPS-HS sample (t=24h). After hexanediol was added, samples were incubated for 24 hours and ThT fluorescence was recorded. **B**. Initial turbidity and final ThT fluorescence of LLPS-HS samples mixed with varying concentration of 1,6-hexanediol. In both A and B, 100 µM tau187 and 3.0 M NaCl were used.

**Figure 4-figure supplement 1.**
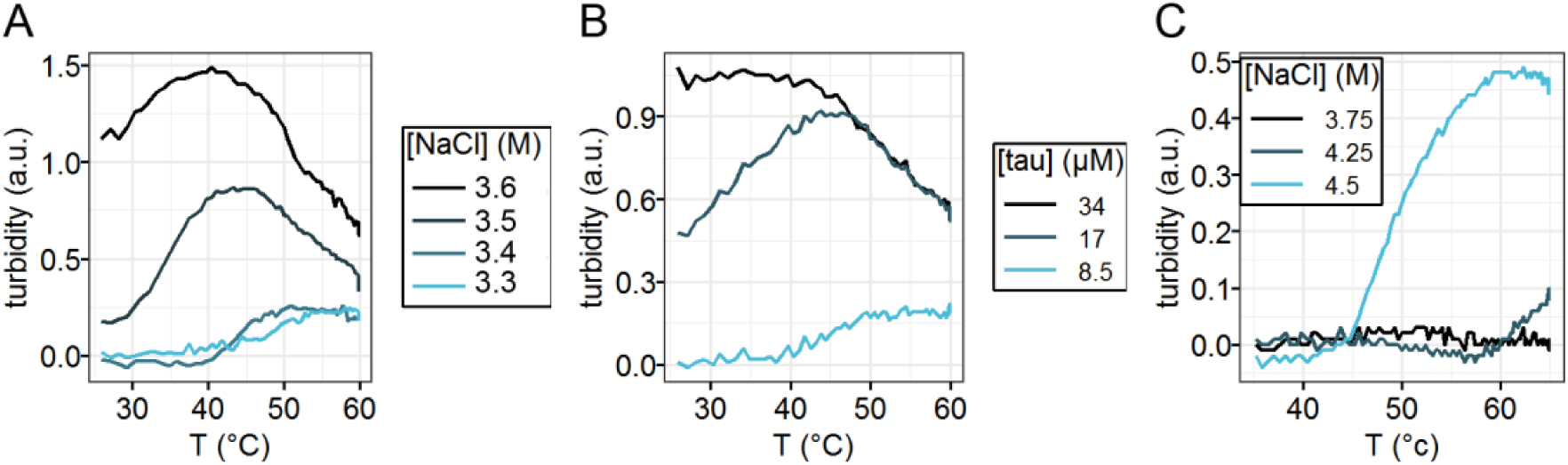
LCST behavior of tau LLPS-high salt. **A-B**. turbidity of tau187 LLPS-high salt upon heating at various [NaCl] with 34 μM tau (A) and at varying [tau] with 3.9 M NaCl (B). **C**. Turbidity of 2N4R (C291SC322S) LLPS-high salt upon heating and cooling at various [NaCl]. 20 μM tau was used. In A-C, samples were prepared at room temperature (25 °C) and heated to 65 °C at a rate of ∼1°C/min.

**Figure 4-figure supplement 2.**
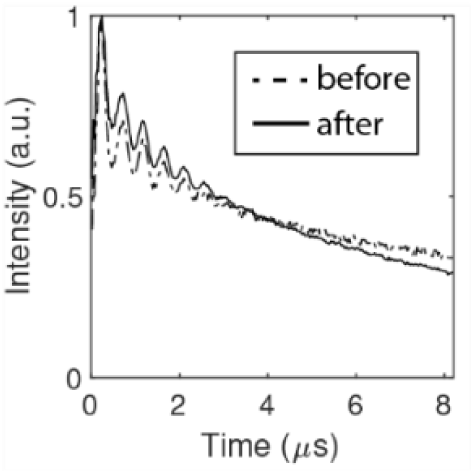
3p-ESEEM data of tau187 upon LLPS-high salt in Figure 5B. Signals were scaled to 1 for comparison.

**Figure 4-figure supplement 3.**
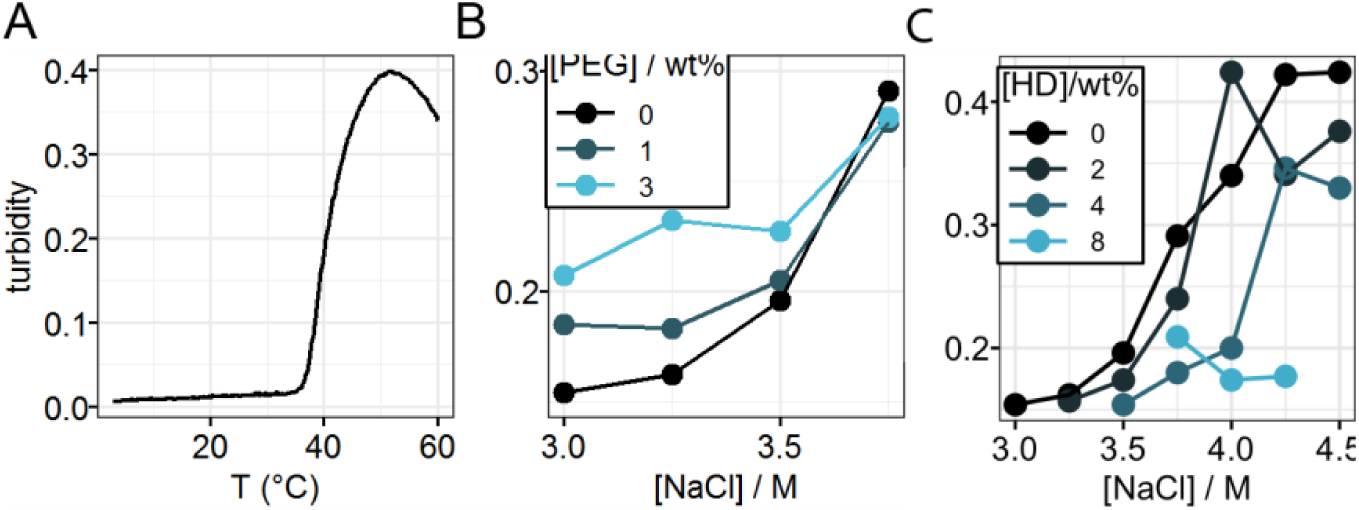
Hydrophobic interaction drives tau LLPS-high salt. **A** Turbidity of tau LLPS-high salt upon heating. 34 µM tau187 and 3.5 M NaCl was used. Samples were pre-cooled to 4 °C and heated up to 65 °C at a rate of 4 °C/min. **B** Representative turbidity vs [NaCl] of tau LLPS-high salt at varying [PEG]. **C**. Turbidity of tau LLPS-high salt at varying [NaCl] and [HD]. In both B and C, 20 µM tau187 was used.

**Figure 4-source data 1.**
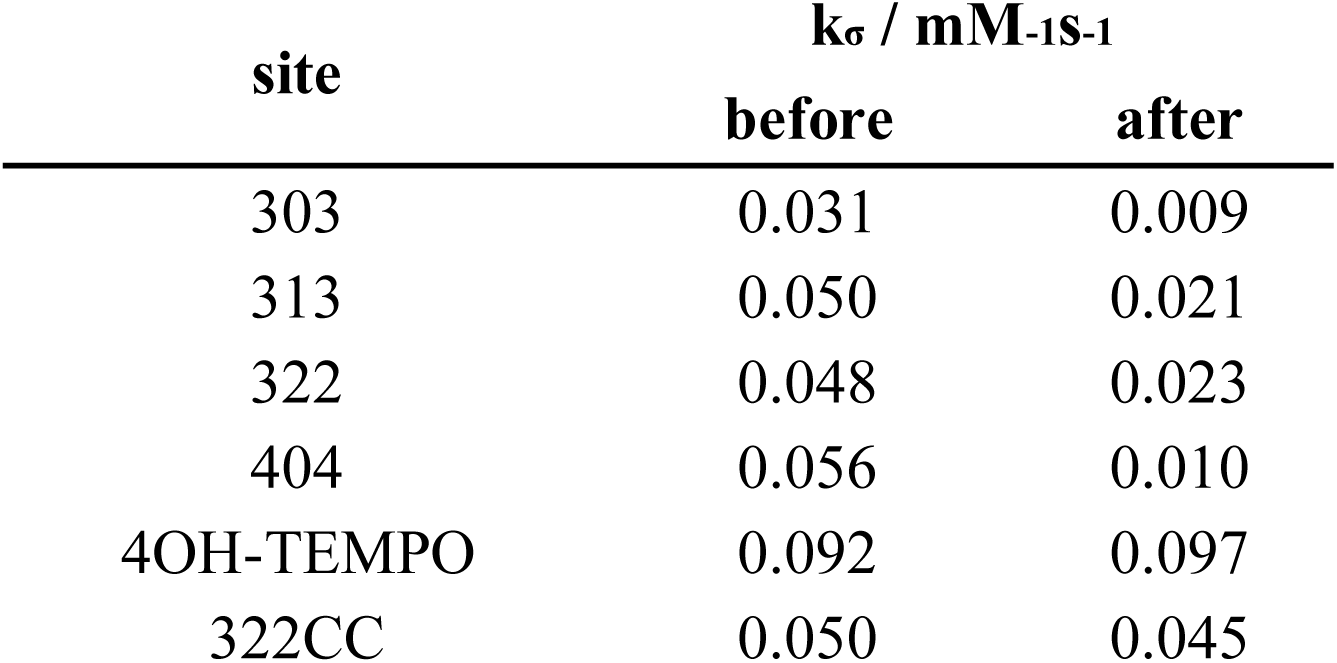
ODNP measured k_σ_ in Figure 4.

**Figure 5-figure supplement 1.**
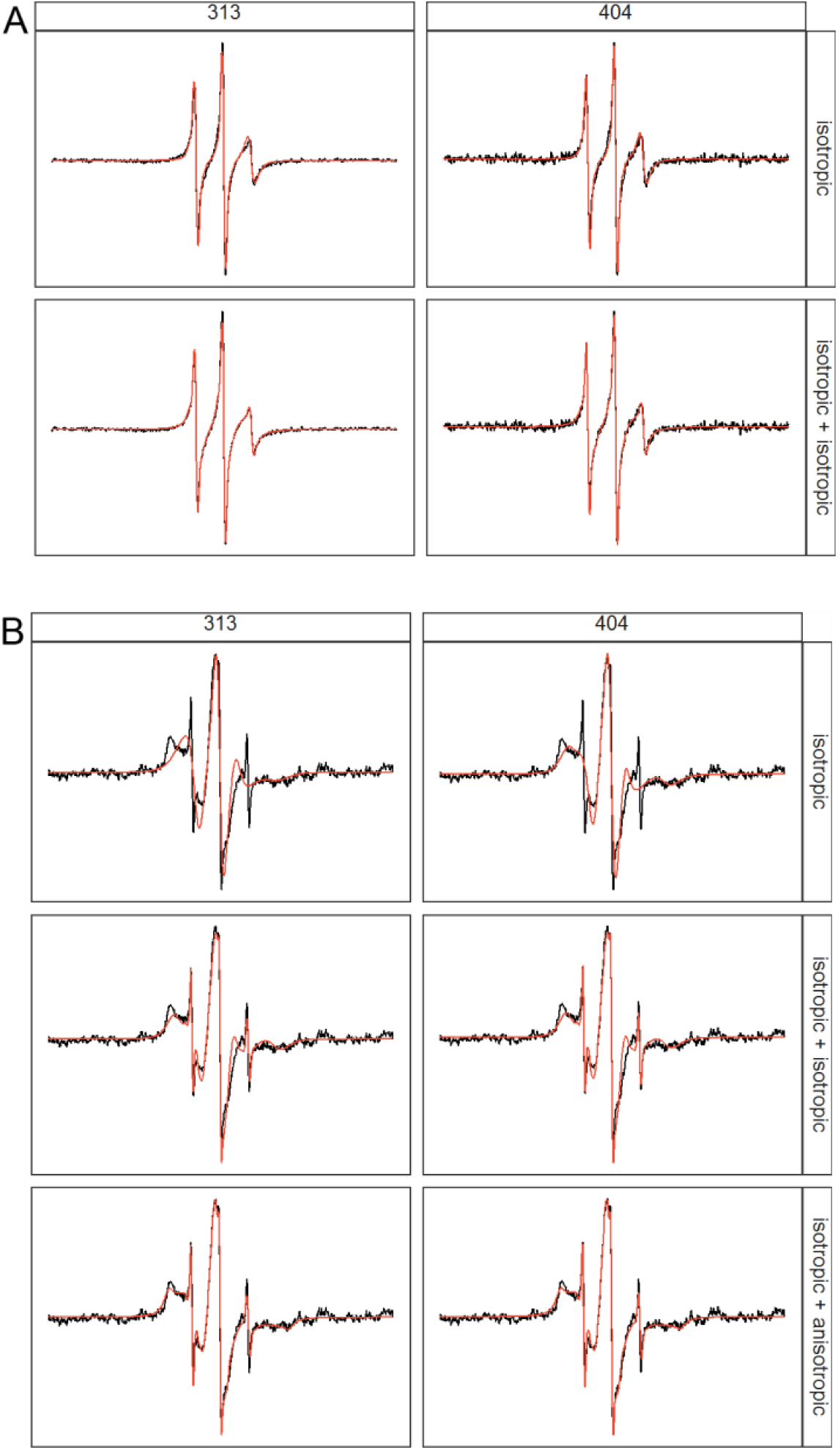
Multicomponent and Anisotropic fitting of cwEPR lineshapes. **A**. Comparison of 1 isotropic component fitting vs 2 isotropic component fitting of tau187 at site 313 and 404 at intermediate [NaCl] of 3.0 M. **B**. Comparison of 1 isotropic, isotropic+isotropic and isotropic+anisotropic components fit of tau187 at site 313 and 404 at very high [NaCl] of 4.5 M.

**Figure 5-figure supplement 2.**
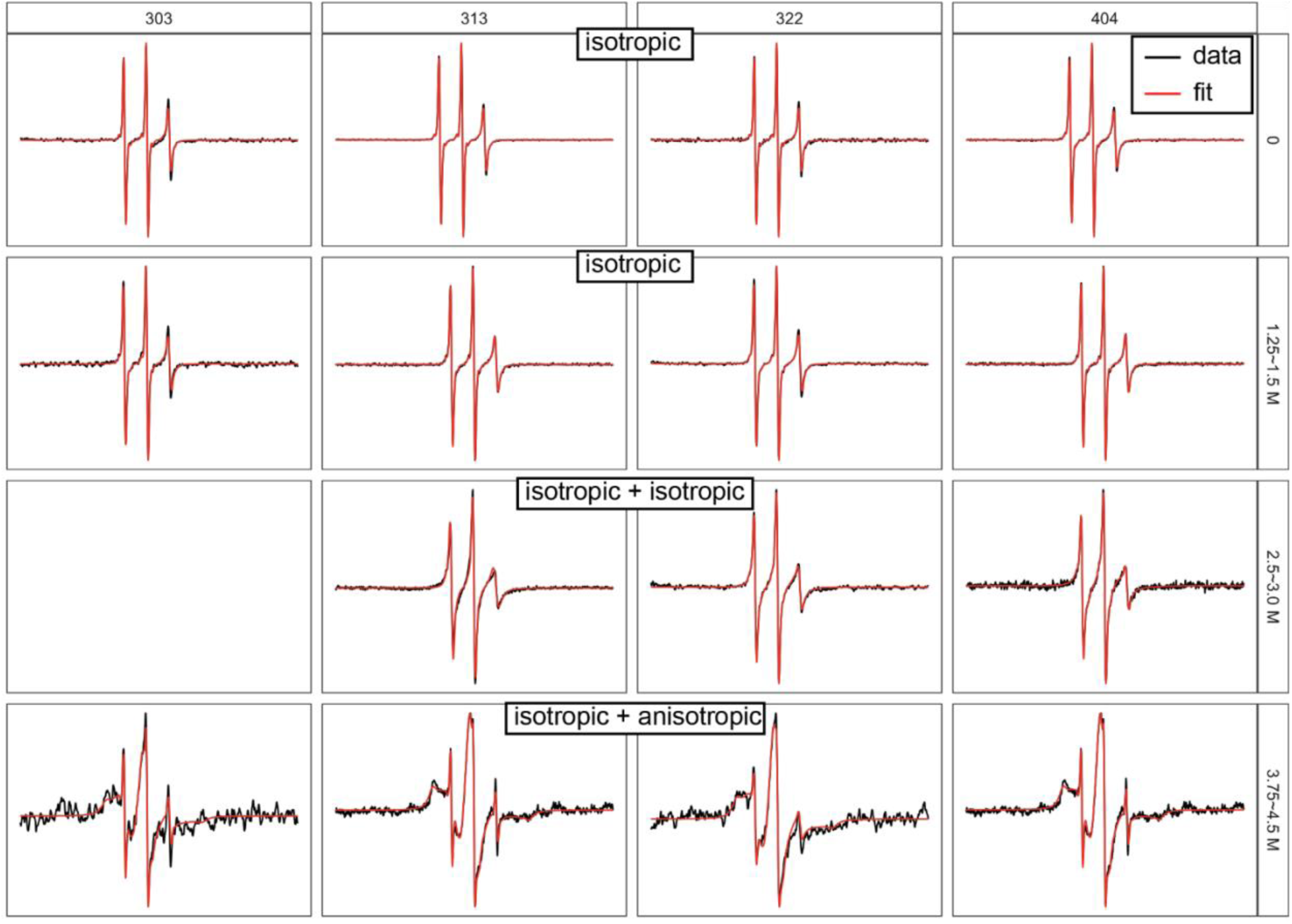
Best fit of cwEPR spectra of tau187 at varying sites and varying [NaCl]. Experiment details and fit results were shown in Figure 5B.

**Figure 5-figure supplement 3.**
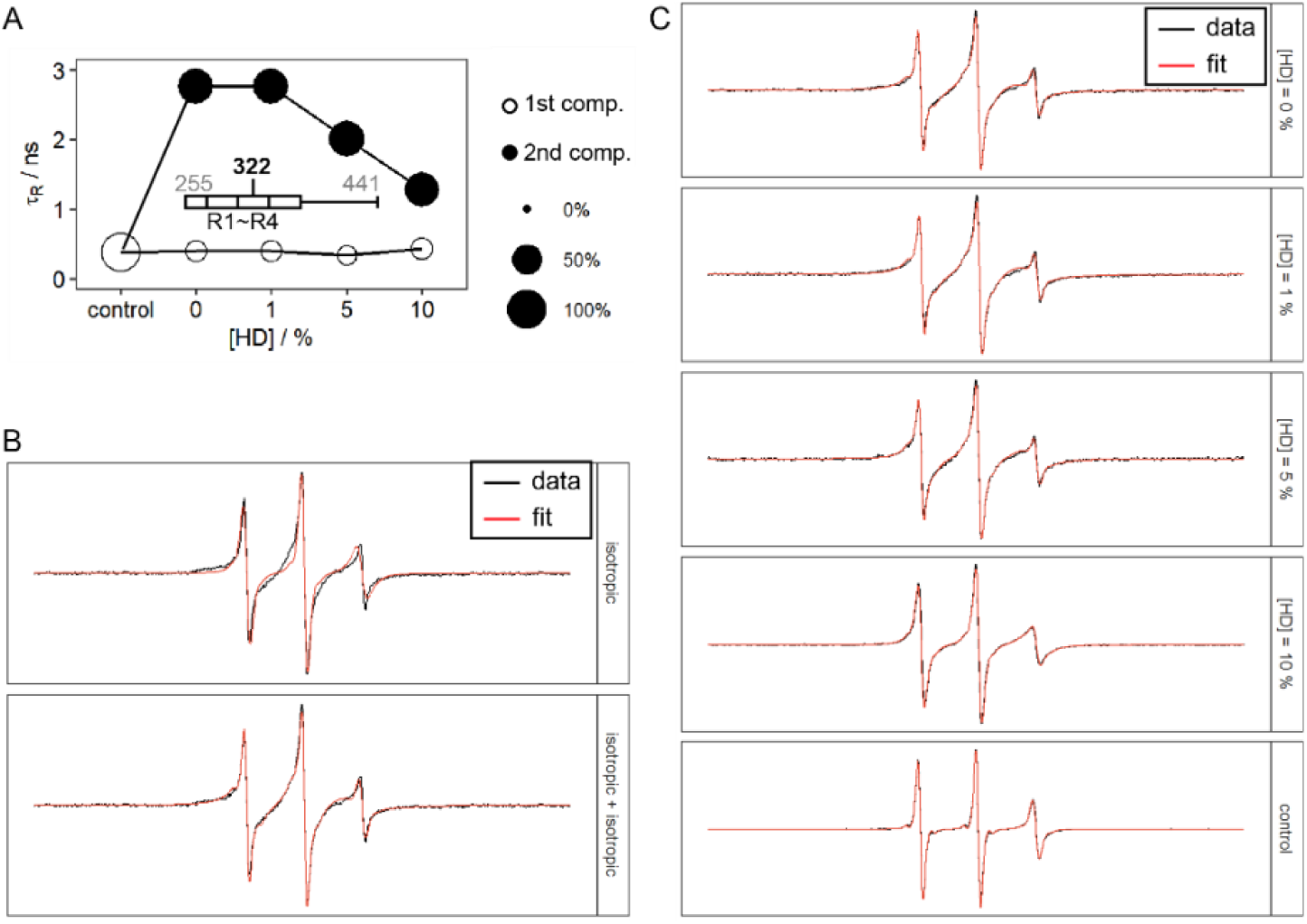
Effects of 1,6-hexanediol on tau LLPS-high salt. **A**. Rotational correlation time of tau LLPS-high salt at varying [HD]. 100 μM tau187P301LC322SL and 2.5 M NaCl were used. **B**. Comparison of fitting cwEPR lineshape using one component vs two component. [NaCl] = 2.5 M, [HD] = 0. **C**. cwEPR lineshape and fitting results of data in A.

**Figure 5-figure supplement 4.**
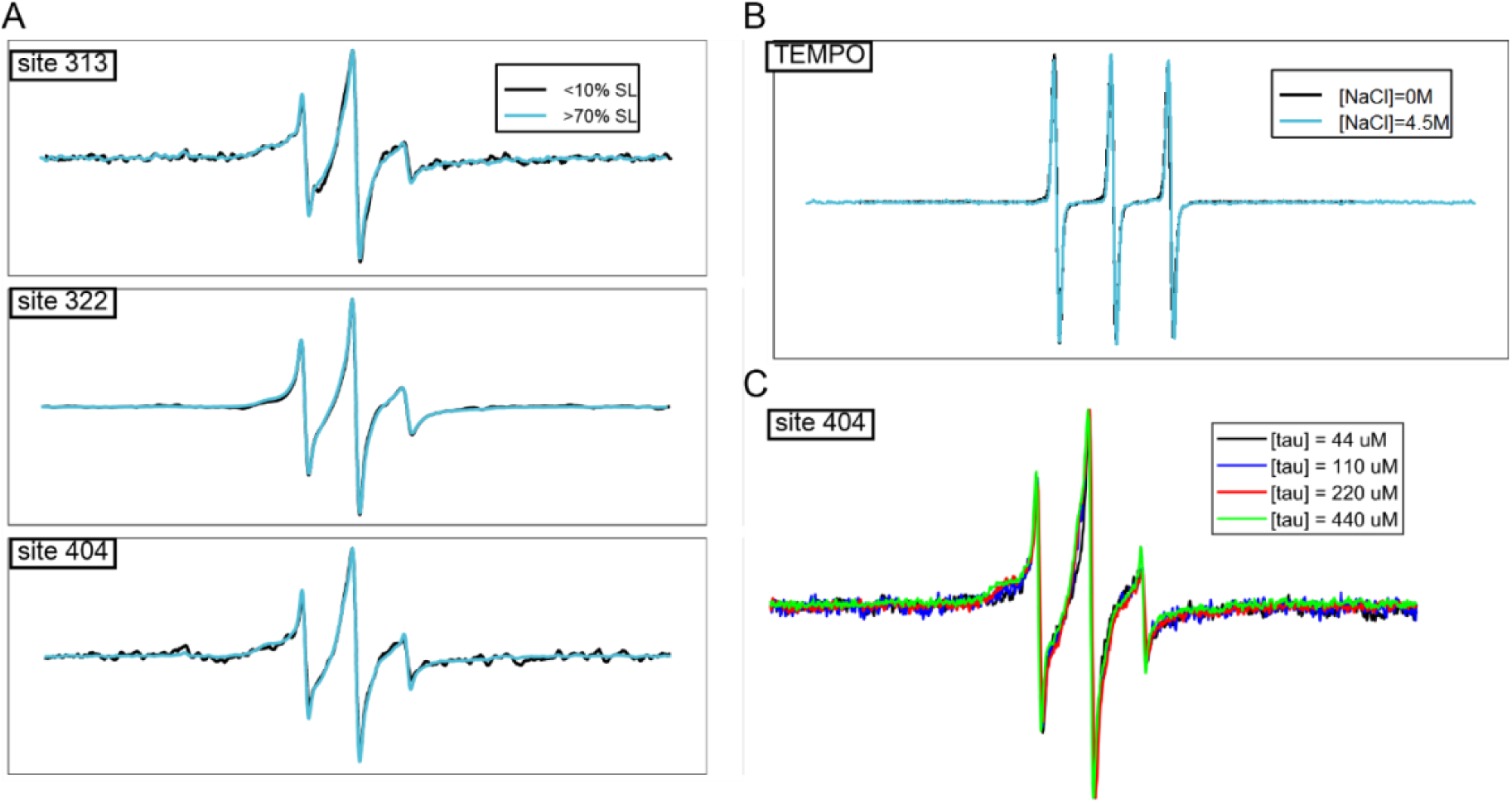
Spin dilution and concentration dependence of cwEPR lineshape. In order to gain information on the oligomer structure, we investigated the site-specific intermolecular interactions by cwEPR. Spin dilution, which consists in mixing 1 spin-labelled tau for 9 non-labeled tau, allows to diminish spin-spin dipolar coupling that could induce EPR lineshape broadening. By comparing cwEPR lineshape of spin-diluted sample vs non-spin-diluted sample, we can determine the population of spins that directly interact within a radius of ∼1.5 nm. **A**. Effects of spin dilution on cwEPR lineshape of tau187 at site 313, 322 and 404. 3.0 M NaCl was used. **B**. Effects of 4.5 M NaCl on 4OH-TEMPO cwEPR lineshape. **C**. Effects of concentrations on cwEPR lineshape of tau187 at site 404 at LLPS-HS conditions with 3.0 M NaCl.

**Figure 5-source data 1.**
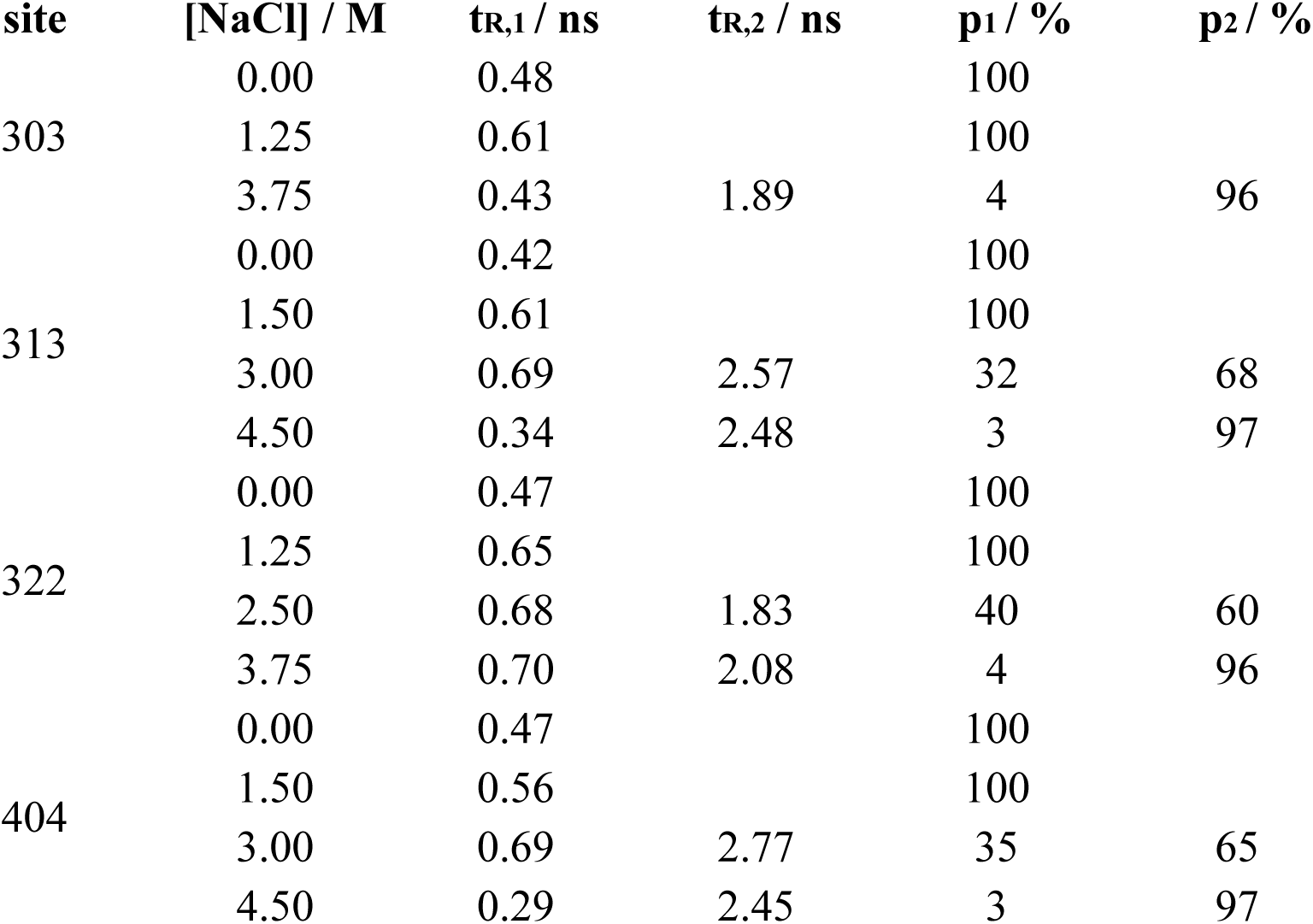
Calculated parameters for cwEPR spectral simulation of tau187 mutants 10 minutes after LLPS at various [NaCl]. Experiment details and fit results were shown in Figure 5B.

**Figure 6-figure supplement 1.**
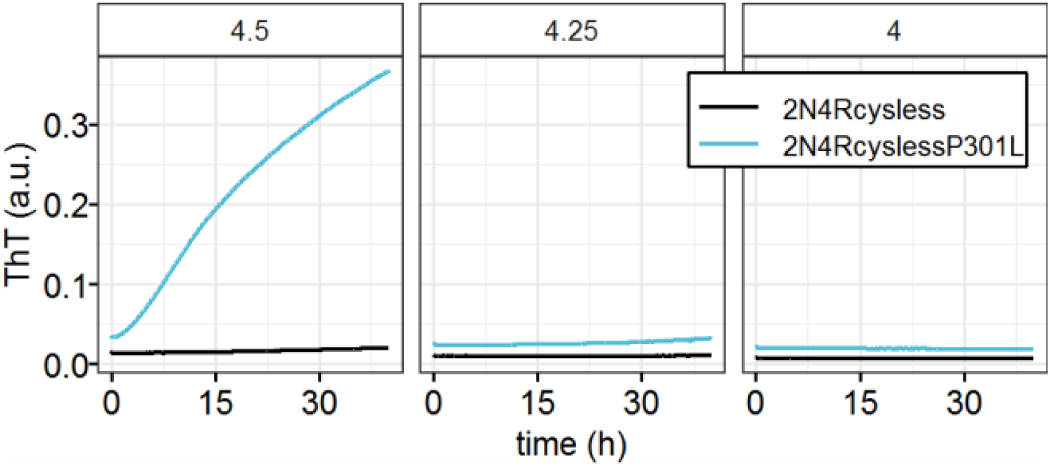
Effects of P301L mutation on amyloid aggregation of 2N4R at high salt concentration. 20 μM 2N4R was used. [NaCl] was shown in panel (unit: M). Samples were incubated at room temperature.

**Figure 6-figure supplement 2.**
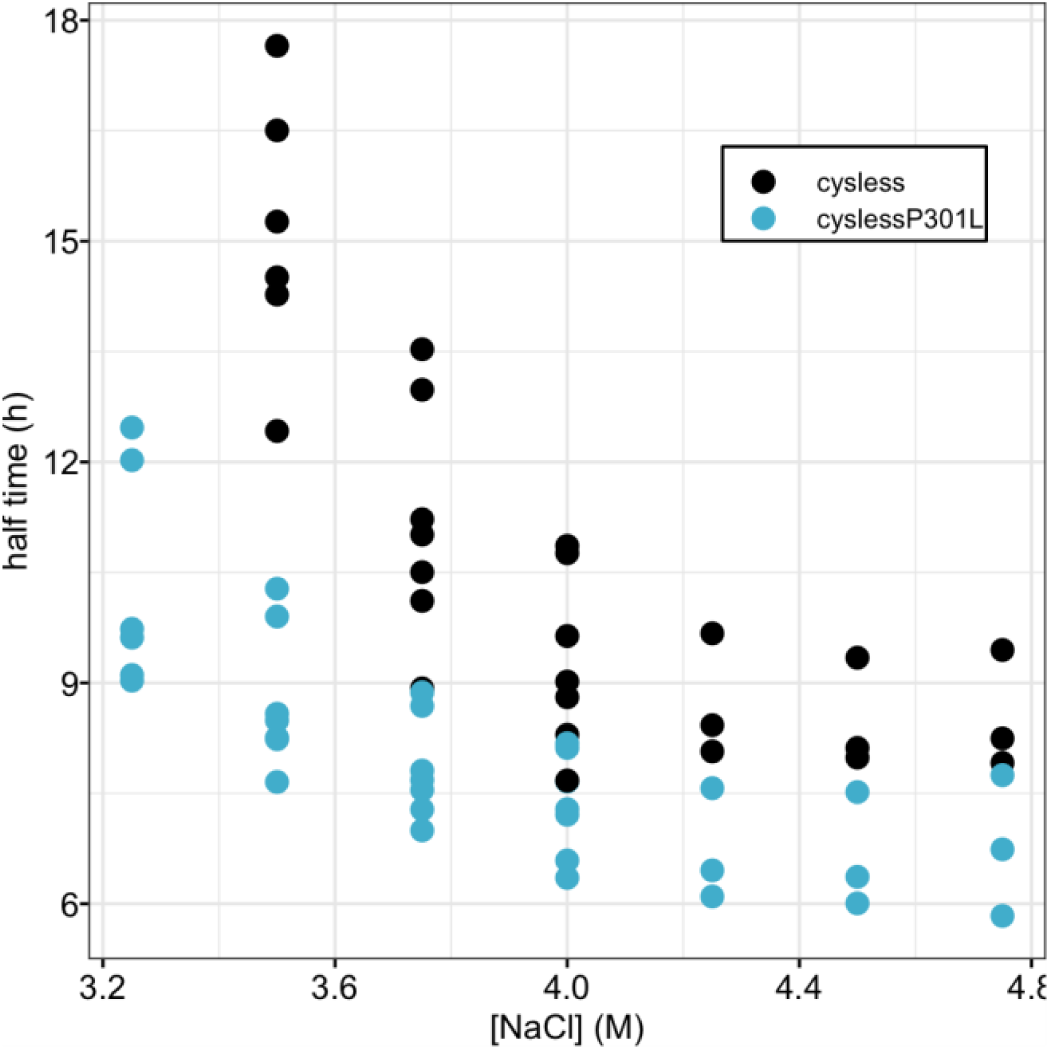
Effects of P301L mutation on half time of ThT fluorescence in high salt droplet conditions. ThT fluorescence readings were recorded at varying [NaCl] using 34 µM tau187C291SC322S (cysless) and tau187C291SC322SP301L (cyslessP301L). Fluorescence readings were fit using a sigmoid function to extract the half time, defined as the time when ThT fluorescence reach half maximum. Data were obtained using n > 5 biological independent repeats.

## Footnotes

1 Estimated from tubulin concentration of 40 µM [5], total tau concentration (bound and free) of 1 ∼ 2 µM [6], dissociation constant of 0.1 ∼ 1 µM [4] as well as stoichiometry of ∼0.5 tau/tubulin dimer [7]. See Materials and Methods for calculation.

